# Ezh2 mediates epigenetic regulation of osteoclastogenesis and bone remodeling in mice

**DOI:** 10.1101/2021.03.24.436787

**Authors:** Jin-Ran Chen, Oxana P. Lazarenko, Dongzheng Gai, Can Li, Michael L. Blackburn, Fenghuang Zhan

**Author notes:** Corresponding author: Jin-Ran Chen, MD, PhD, Arkansas Children’s Nutrition Center, Slot 512-20B, 15 Children’s Way, Little Rock, AR 72202.

## Abstract

Osteoclasts derived from hematopoietic stem cells control bone resorption. Identifying novel molecules that can epigenetically regulate osteoclastogenesis has been an important basic and clinical issue. The polycomb group (PcG) protein enhancer of zeste homologue 2 (Ezh2), a histone lysine methyltransferase is associated with epigenetic regulation of numerous cellular processes, it is not yet clear on its involvement in bone cell development and homeostasis. Here, we crossed LysM-Cre mice with Ezh2^flox/flox^ mice to delete Ezh2 in myeloid cell lineage mature macrophages. Conditional deletion of Ezh2 in macrophages resulted in significant increases in postnatal bone growth in the first 6 months of life, but tibia length and body weight gains were not different in knockout mice compared with their wild type controls. Significantly decreased osteoclastogenesis but increased bone mass without osteopetrosis were found in Ezh2 conditional knockout (CKO) mice. In contrast to female mice, one floxed Ezh2 gene copy recombinant with LysM-Cre^+^ (Ezh2^flox/+^LysM-Cre^+^) produced increased bone mass in young adult male mice compared with control mice (Ezh2^flox/flox^, LysM-Cre^+^ and wild type). Inflammatory milieu in bone was significantly lower in both male and female CKO mice compared with their respective controls. Deletion of Ezh2 in macrophages triggered increased gene expression of osteoclast suppressors, IRF8, MafB and Arg1 due to decreased Ezh2-induced trimethylation of H3K27me3. Conversely, NFATc1 and Cathepsin k expression were decreased. These findings suggest that pre-osteoclastic cell differentiation is under epigenetic control of osteoclast suppressive gene expression via an Ezh2-dependent mechanisms.

## Introduction

Osteoclasts derived from hematopoietic stem cells and during early development, monitor osteoblasts to control bone resorption and to shape newly formed bone (Siddiqui & Partridge, 2016; Bethel *et al*, 2011; Zaidi *et al*, 2018). While osteoclastic bone resorption is an important cellular function in the development and physiology of the skeleton, suppressing osteoclastogenesis is considered an effective therapeutic approach for bone destructive or resorptive diseases, such as osteoporosis and rheumatoid arthritis (Hirayama *et al*, 2002; Tanaka, 2018). On the other hand, defective osteoclastic bone resorption leads to accumulation of bone with altered architecture, resulting in osteopetrotic bone susceptible to fractures (Stattin *et al*, 2017). Receptor activator of NF-κB ligand (RANKL) and macrophage colony-stimulating factor (M-CSF) (Feng & Teitelbaum, 2013) have been characterized as two crucial factors that influence the formations of osteoclasts. Moreover, it is also known that many inflammatory factors such as TNF-α mediates osteoclast formation, although the role of those inflammatory factors including TNF-α signaling in osteoclastogenesis remains poorly understood (Cao et al, 2017). Myeloid hematopoietic cells are thought to be precursors for osteoclasts, other cell lineages derived from hematopoietic stem cells share similar differentiation signaling pathways, it is not yet clear how to target specific osteoclastogenic signaling for intervening persistently increased osteoclastic cell differentiation and bone resorption. Therefore, identifying novel molecules that can regulate, especially epigenetically control osteoclastogenesis has been an important basic and clinical issue.

As pathologies of osteoporosis or rheumatoid arthritis are not characterized by common targetable genetic alterations, epigenetic mechanisms and strategies for preventions might be relevant and promising options. Indeed, epigenetic regulation of osteoclastic cell differentiation and activity has been increasingly investigated (Yasui *et al*, 2011; Kyung-Hyun, 2019). Recent cell culture studies suggested that generalized accumulation of methylation at the CpG island in the mouse RANKL promoter may result in the recruitment of histone deacetylase and chromatin condensation, leading to the stable epigenetic silencing of RANKL (Kitazawa & Kitazawa, 2007). Moreover, DNA methyltransferase 3a (DNMT3a) has been reported an important function in promoting osteoclastogenesis by increasing DNA methylation of anti-osteoclastogenic genes (Nishikawa *et al*, 2015). DNA methylation in the NFATc1 has profound effects on osteoclast differentiation and activity (Kurotaki *et al*, 2020; Yasui *et al*, 2011). These studies are somewhat limited in mainly being performed *in vitro,* and often focus on a few genes without determining upstream factors (Akimzhanov *et al*, 2008). Furthermore, DNA methylation is not an isolated event, but also affects and is itself affected by other gene regulatory mechanisms, such as gene silencing via histone modification (methylation or acetylation). While there is a dearth of evidence on epigenetic regulation of bone cell development in association with changes of life styles during early growth, we have previously hypothesized that high fat diet (HFD)-induced obesity may alter enhancer of zeste (Ezh) polycomb repressive complex subunit expression to epigenetically control osteoclastogenesis and therefore bone resorption (Chen *et al*, 2018). Such regulation may be through activation of Ezh2 to promote osteoclastogenesis by downregulating IRF8, a negative regulator of osteoclastogenesis (Fang *et al*, 2016).

Ezh2, a histone lysine methyltransferase associated with transcriptional repression (Donaldson-Collier *et al*, 2019), catalyzes the addition of methyl groups to histone H3 at Lys 27 (H3K27me3) in target genes involved in numerous cellular processes, predominantly leading to gene repression (Lavarone *et al*, 2019). Loss or inhibition of Ezh2 leads to enhanced osteogenic but inhibited adipogenic differentiation of bone marrow-derived mesenchymal stem cells (BMSCs) both from rodents and humans (Hemming *et al*, 2017), as well as in murine pre-osteoblastic cell line MC3T3 (Dudakovic *et al*, 2020). Expression of Ezh2 suppresses the osteogenic genes and ligand-dependent signaling pathways (e.g., WNT, PTH, and BMP2) to favor adipogenic differentiation (Wang *et al*, 2010). Several studies have shown that inhibition of Ezh2 prevents estrogen deficiency-induced bone loss in animal models (Dudakovic *et al*, 2016). Despite evidence for a role of Ezh2 in bone cell differentiation, it is not known if this system is involved in maternal HFD-obesity-associated offspring programming of bone development. Depend on different Cre mouse models were used, deletion of Ezh2 in osteoblastic cell lineage resulted in unclear bone developmental patterns (Chen *et al*, 2020). We recently reported that, using Osterix Cre mouse model, although deletion of Ezh2 in osteoblast precursors resulted in clear osteogenic gene higher expression in bone, bone mass evaluated by peripheral quantitative CT scan showed obscure phenotype when conditional Ezh2 deletion mice compared with Osterix Cre or Ezh^flox/flox^ mice as controls (Levaot *et al*, 2015). Bone development and remodeling are not separated osteoblastic events, two other types of bone cells, osteoclasts and osteocytes are also involved. Our interests in determining the role of Ezh2 during osteoclast differentiation and bone resorption derived from the evidence that Ezh2 might be involved in bone osteoblastic cell activity and differentiation.

In the present report, we set to examine if deletion of Ezh2 gene in osteoclast precursors would result in changes of bone mass phenotype in young and adult mice, and what is the pattern of post-natal skeletal development after deletion of Ezh2 gene in osteoclast precursors. In addition, we present if Ezh2 gene expression in pre-osteoclasts plays different roles in bone homeostasis in different sex of mice, and inform if Ezh2 gene controls or modifies specific osteoclast gene expression to epigenetically regulate bone resorption.

## Results

### Changes of Ezh2 and osteoclast suppressive gene expression during bone cell and post-natal bone development

We determined Ezh2 and osteoclast suppressive gene expressions during bone development of wild type C57Bl mice post weaning (30 days old) to 6 months old, and during osteoclast and osteoblast differentiation. We found that, although Ezh2 mRNA was abundantly expressed in bone (total RNA was isolated from spine L4), there were no significant differences among those male wild type mice of 30 days old, 56 days old and 6 months old groups (Fig 1A). Expression of RIF8, a key osteoclast suppressive gene, was highest in bone from 56 days old mouse group compared with two other age groups (Fig 1B). However, the osteoblast bone formation marker osteocalcin (Fig 1C) and osteoclast bone resorption marker Cathepsin K (Fig 1D) mRNA expression were both significantly lower in 6 months old group compared with either 30 or 56 days old mouse group, indicating bone remodeling in 6 months old young adult mice is lower than in younger mice. The protein expression of Ezh2 (Fig 1E) in bone was lower in mice 30 days old compared with those of 56 days and 6 months old mice, but IRF8 and Arg1 (another osteoclast suppressive factor) expression were significantly lower in 6 month old mice compared with those of either 30 days or 56 days old mouse groups (Fig 1E). On the other hand, NFATc1 (a positive osteoclastogenic regulator) protein expression was significantly higher in 6 month old mice compared with two other age groups, indicating osteoclastogenesis increase with age (regardless of lower bone remodeling). Since RNA and protein isolated from L4 bone were mixed with other cell types, we took murine osteoclastic and osteoblastic cell lines to investigate those gene expressions during bone cell differentiation. In the presence of RANKL, it took about 5 days to differentiate osteoclast precursors to mature multi-nuclear mature osteoclasts marine cell line RAW264.7 osteoclastic cell cultures (Fig 1F). During osteoclast differentiation, Ezh2 gene expression significantly dropped from day 3 to day 5 (Fig 1G) compared with its expression in osteoclastic cells on day 1. On the other hand, mRNA expression of IRF8 and MafB (another negative osteoclast regulator), two osteoclast inhibitory genes, were significantly increased at day 3 and day 5 compared with their expression in day 1 (Fig 1G), indicating Ezh2 and osteoclast inhibitory gene expression has inverse association. In osteoblast stromal cell line ST2 cell cultures, it took 7 days for ST2 cells to differentiate into ALP positive mature osteoblast in the presence of osteoblast culture medium (Fig 1H). During this period, Ezh2 mRNA expression was significantly lower on day 3 and 7 compared to day 1 (Fig 1I). Bone specific ALP mRNA expression was significantly increased on day 3 and 7 compared with on day 1, indicating Ezh2 and ALP expression has an inverse relationship during osteoblast differentiation. However, osteocalcin mRNA expression was significantly lower on day 3 and 7 compared with its expression on day 1 (Fig 1I). These data indicated that Ezh2 expression is higher during both osteoclast and osteoblast early differentiation compared with their later differentiation stages, and Ezh2 is inversely associated with either osteoclast suppressive gene or osteoblastic marker gene expression.

**Figure 1.**
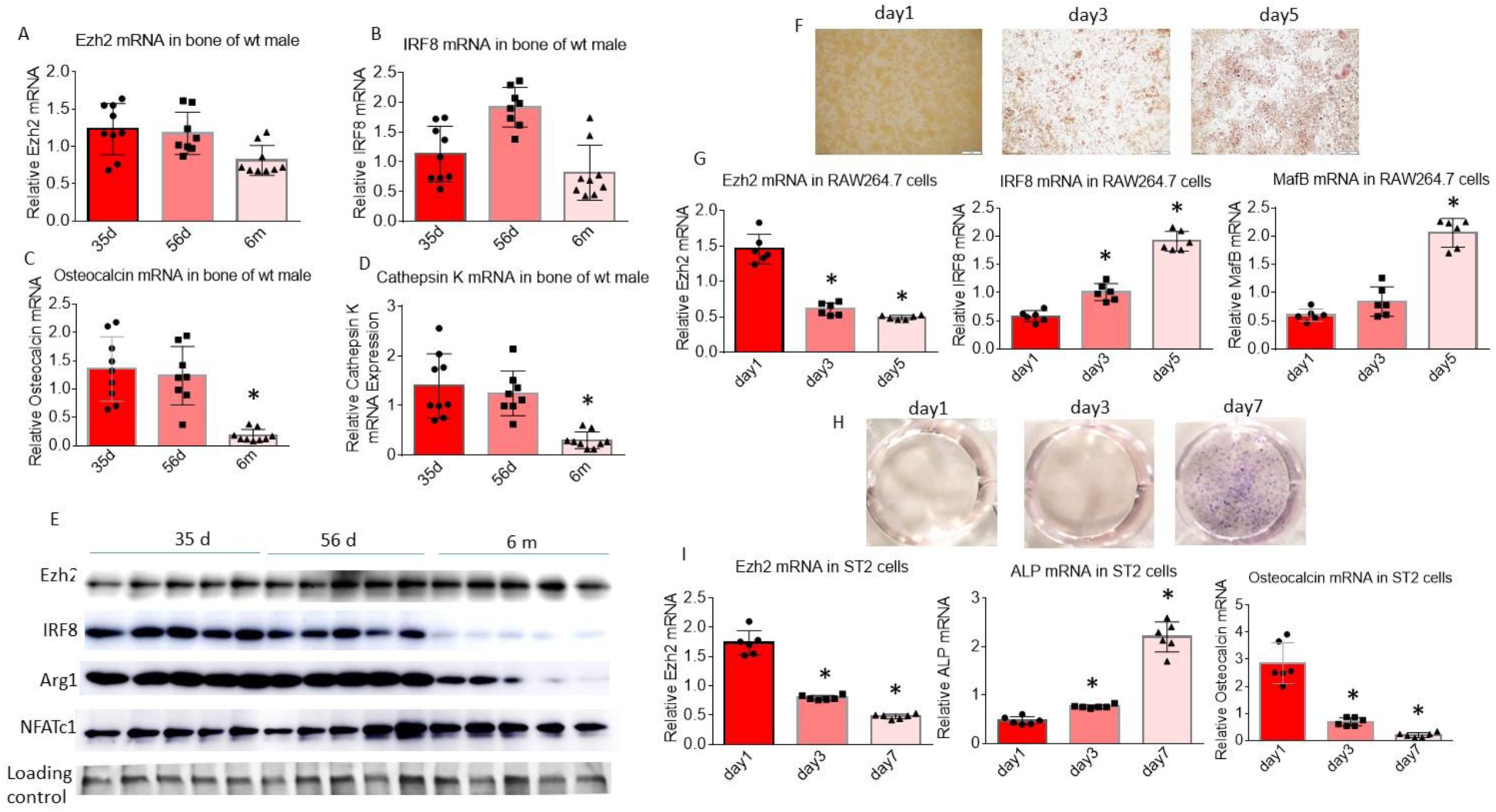
Ezh2 expression in bone and in bone cells during their differentiation. Total RNA was isolated from spine L4 of male 30-days-old, 56-days-old and 6-months-old wild type mice, real-time PCR determined Ezh2 **(A)**, IRF8 **(B)**, osteocalcin **(C)** and Cathepsin K **(D)** mRNA expression. **(E)** Total protein was isolated from spine L3 of male 30-days-old, 56-days-old and 6-months-old wild type mice, Western blots determined Ezh2, IRF8, Arg1 and NFATc1 protein expression, samples were pooled 5 per group to load to mini-gels. **(F)** Representative cell culture pictures of TRAPase staining of RAW264.7 cells in the presence of 30 ng/ml of RANKL on day 1, day 3 and day 5. **(G)** Real-time PCR determined Ezh2, IRF8 and MafB mRNA expression in RAW264.7 cells cultured in presence of 30 ng/ml of RANKL on day 1, day 3 and day 5, six culture wells of 12-well culture plates per group. **(H)** Representative cell culture pictures of ALP staining of ST2 cells in the osteoblast differentiation medium on day 1, day 3 and day 7. (I) Real-time PCR determined Ezh2, ALP and osteocalcin mRNA expression in ST2 cells cultured in osteoblast differentiation medium on day 1, day 3 and day 7, six culture wells of 12-well culture plates per group. * p<0.05 by one-way ANOVA followed by Tukey’s post hoc test.

### Ezh2 gene expression in osteoclastic cells is not required for maintaining better bone mass in 6 months old young adult mice

The inverse association between Ezh2 and osteoclast inhibitory gene expression during osteoclast differentiation triggered us to investigate the role of Ezh2 gene on controlling osteoclastogensis *in vivo* and postnatal bone development. We have used a large amount of mouse numbers and created substantial amount of data to determine postnatal bone mass in osteoclastic cell specific Ezh2 gene knockout mice. Bone structures were assessed using micro-CT, we first show those results in 6 months old male and female mice. As shown in representative scans of tibia of male mice in Fig 2A, among parameters which were analyzed on trabecular bone, we present percentage of trabecular bone volume BV/TV (bone volume / total tissue volume), actual bone volume (BV), bone surface (BS), BS/TV, trabecular number (Tb.N), trabecular separation (Tb.Sp) from Ezh2^flox/flox^/LysM-Cre^+^ (CKO, conditional knockout), Ezh2^flox/flox^ (f/f, flox control), LysM-Cre^+^ (Cre^+^, Cre^+^ control), Ezh2^flox/+^/LysM-Cre^+^ (f/+ Cre^+^, Ezh2 one copy floxed with LysM-Cre positive), Ezh2^flox/+^/LysM-Cre^-^ (f/+ Cre^-^, Ezh2 one copy floxed with LysM-Cre negative) and wild type (Wt) control mice in Fig 1B. One-way analysis of variance (ANOVA) followed by Tukey’s multiple comparison analysis was performed, there was no significant difference of BV/TV between CKO and f/+ Cre^+^ mouse groups, but it was significantly higher in those two groups compared with any other four different genotypic control groups of mice (f/f, Cre^+^, f/+ Cre^-^ and Wt) (Fig 2B). There were no significant differences among those four different genotype control groups of mice on BV/TV. Like BV/TV, other parameters of BV, BS, BS/TV and Tb.N showed similar patterns among CKO and other four groups (Fig 2B). In contrast, Tb.Sp was significantly lower in CKO and f/+ Cre^+^ mouse groups compared with all other genotypes (Fig 2B). In females, as shown in representative scans of tibia from all genotypic mouse groups in Fig 2C, we found BV/TV was significantly higher in CKO group compared to other genotypic and wild type mouse groups (Fig 2D). It was not like we observed in males that there were no differences between CKO and f/+ Cre+ mouse groups on such as BV/TV, we found that BV, BS, BS/TV and Tb.N were all significantly higher, but Tb.Sp was significantly lower only in CKO mice compared with those from any other genotypic and wild type mouse groups (Fig 2D). Micro-CT was also performed on vertebrae (L5), as shown in representative scans of L5 of male mice in Fig 3A, BV/TV was significantly higher in CKO and f/+ Cre+ mouse groups compared with any other four genotypic mouse groups (Fig 3B), but BV/TV was also significantly higher in CKO mice compared with f/+ Cre+ mice (Fig 3B). Significant difference of BV was only found between CKO and f/f mice (Fig 3B). BS and BS/TV results were similar to those of the tibia (Fig 3B). Tb.N was significantly higher in CKO compared with all other groups (Fig 3B). Tb.Sp in CKO mice was significantly lower compared with f/f, Cre+, f/+ Cre- and Wt mice, but was not different when compared to f/+ Cre+ mice (Fig 3B). In females, as shown in micro-CT representative L5 scans in Fig 3C, BV/TV in CKO mice had significantly higher than in any other genotypic mouse groups (Fig 3D). BV in CKO and f/+ Cre+ mice were significantly different than in f/f and f/+ Cre-mice (Fig 3D). BS/TV and Tb.N were significantly higher in CKO mice compared with any other groups (Fig 3D), but BS and Tb.Sp were not significantly different across groups (Fig 3D). We found no differences of mouse body weights and tibia lengths among groups and additional micro-CT data of tibia trabecular of 6 months old male mice in Supplemental Table 1 and female mice in Supplemental Table 2. Moreover, micro-CT tibia cortical parameters were also analyzed, those significantly changed parameters in CKO mice compared with other genotypic mouse groups can be found in Supplemental Table 3 for males and in Supplemental Table 4 for females.

**Figure 2.**
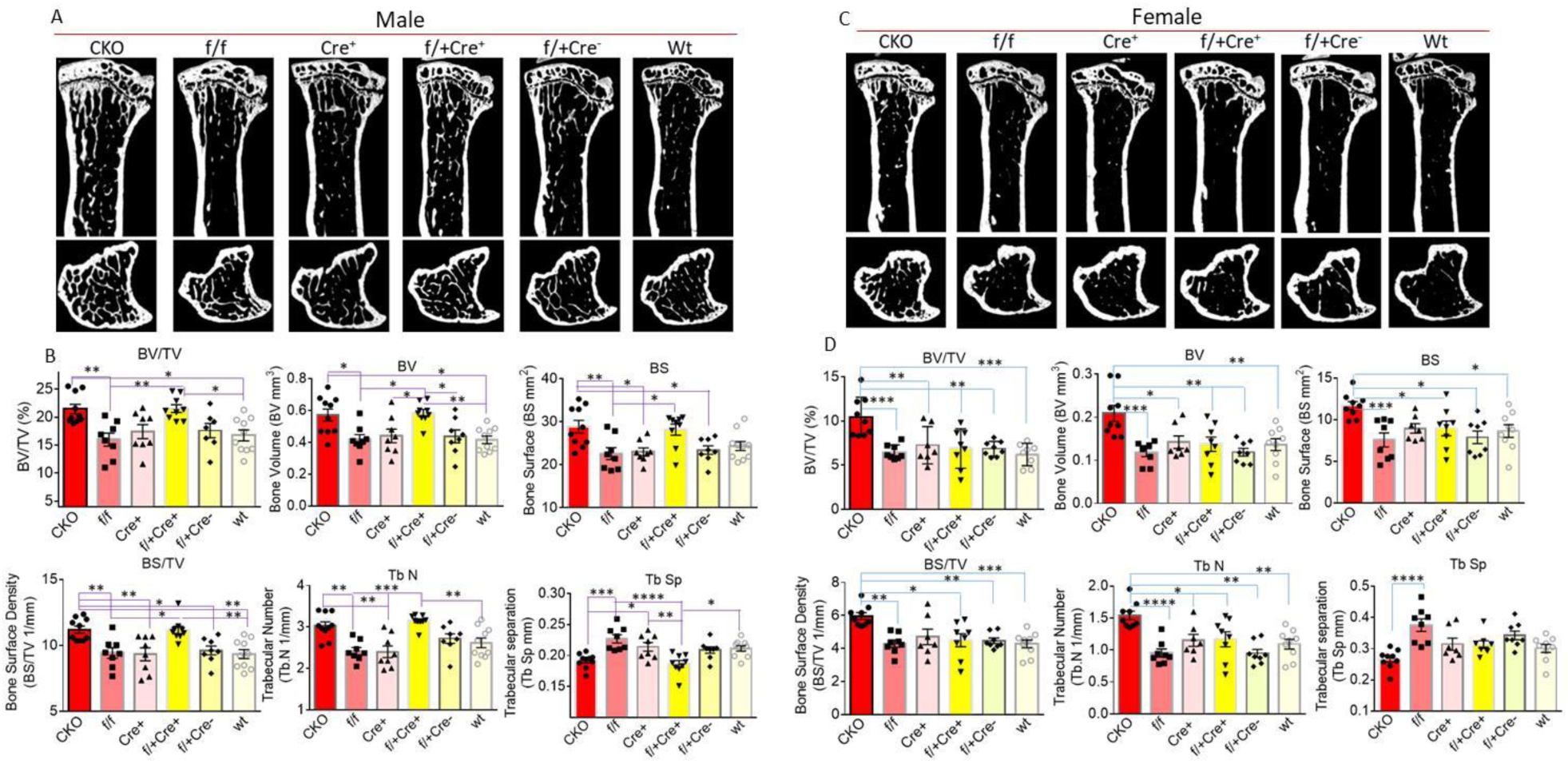
Micro-CT bone phenotype of tibia of 6 months old male and female Ezh2^flox/flox^/LysM-Cre^+^ CKO and other genotypic and wild type control mice. **(A)** Representative micro-CT images of male proximal tibia from one sample from each group of mice, upper panel shows sagittal view and lower panel shows transverse view, white lines and dots indicates trabecular or cortical bone tissues. **(B)** Micro-CT measures of six parameters from trabecular tibias from CKO (Ezh2^flox/flox^/LysM-Cre^+^, n=10), f/f (Ezh2^flox/flox^ control, n=8), Cre^+^ (LysM-Cre^+^ control, n=8), f/+Cre^+^ (Ezh2^flox/+^/LysM-Cre^+^, n=9), f/+Cre^-^ (Ezh2^flox/+^/LysM-Cre^-^, n=7) and Wt (wild type control, n=9) male mice. BV/TV, bone volume/total tissue volume; BV, bone volume; BS, bone surface; BS/TV, bone surface density; Tb.N, trabecular number; Tb.Sp, trabecular separation. **(C)** Representative micro-CT images of female proximal tibia from one sample from each group of mice, upper panel shows sagittal view and lower panel shows transverse view, white lines and dots indicates trabecular or cortical bone tissues. **(D)** Micro-CT measures of six parameters from trabecular tibias from CKO (n=9), f/f (n=8), Cre^+^ (n=7), f/+Cre^+^ (n=8), f/+Cre^-^ (n=8) and Wt (n=8) female mice. Parameter BV/TV, BV, BS, BS/TV, Tb.N and Tb.Sp were presented. Data are expressed as mean ± SD, analyzed by one-way ANOVA, additionally, * p<0.05, ** p<0.01, *** p<0.001, **** p<0.0001 by Tukey’s multiple comparison.

**Figure 3.**
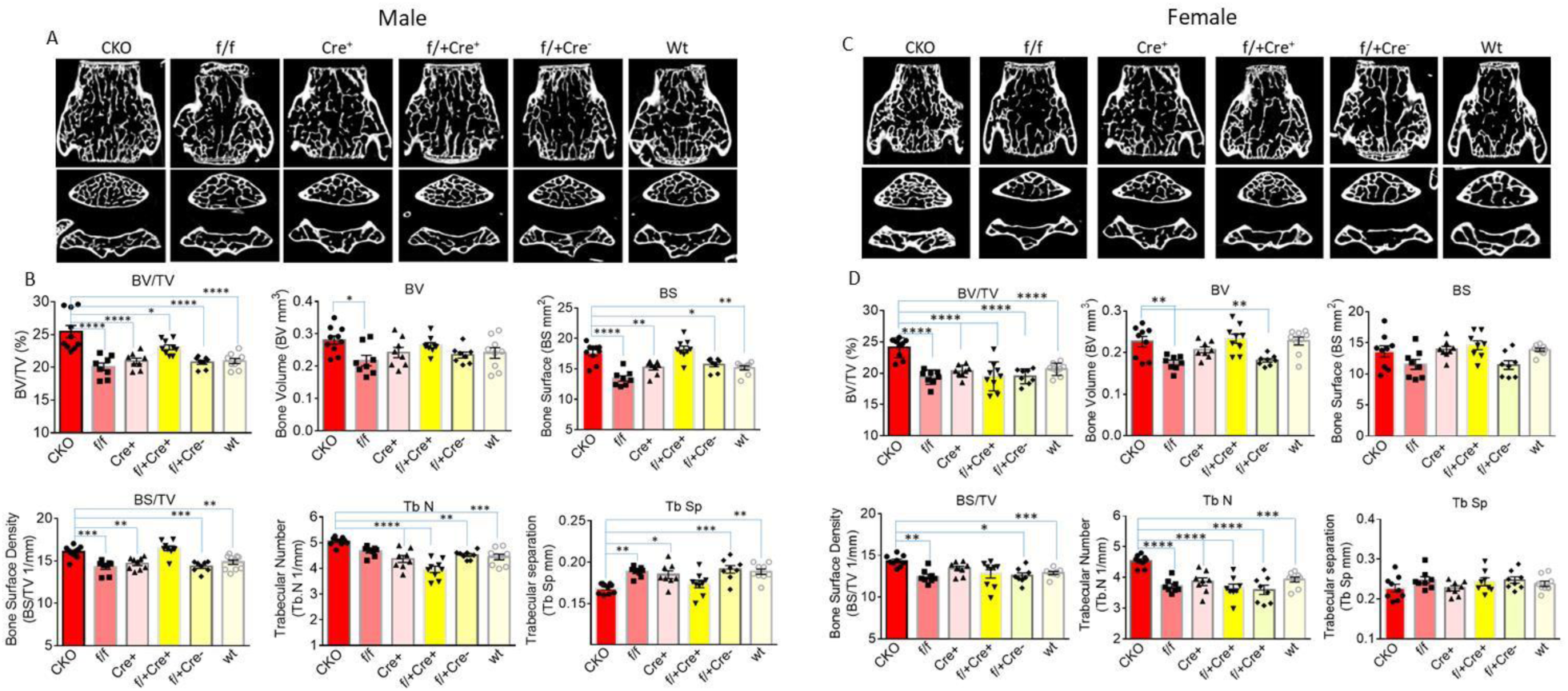
Micro-CT bone phenotype of spine L5 of 6 months old male and female Ezh2^flox/flox^/LysM-Cre^+^ CKO and other genotypic and wild type control mice. **(A)** Representative micro-CT images of male L5 from one sample from each group of mice, upper panel shows sagittal view and lower panel shows transverse view, white lines and dots indicates trabecular or cortical bone tissues. **(B)** Micro-CT measures of six parameters from trabecular L5 from CKO (Ezh2^flox/flox^/LysM-Cre^+^, n=10), f/f (Ezh2^flox/flox^ control, n=8), Cre^+^ (LysM-Cre^+^ control, n=8), f/+Cre^+^ (Ezh2^flox/+^/LysM-Cre^+^, n=9), f/+Cre^-^ (Ezh2^flox/+^/LysM-Cre^-^, n=7) and Wt (wild type control, n=9) male mice. BV/TV, bone volume/total tissue volume; BV, bone volume; BS, bone surface; BS/TV, bone surface density; Tb.N, trabecular number; Tb.Sp, trabecular separation. **(C)** Representative micro-CT images of female L5 from one sample from each group of mice, upper panel shows sagittal view and lower panel shows transverse view, white lines and dots indicates trabecular or cortical bone tissues. **(D)** Micro-CT measures of six parameters from trabecular L5 from CKO (n=9), f/f (n=8), Cre^+^ (n=7), f/+Cre^+^ (n=8), f/+Cre^-^ (n=8) and Wt (n=8) female mice. Parameter BV/TV, BV, BS, BS/TV, Tb.N and Tb.Sp were presented. Data are expressed as mean ± SD, analyzed by one-way ANOVA, additionally, * p<0.05, ** p<0.01, *** p<0.001, **** p<0.0001 by Tukey’s multiple comparison.

### Postnatal bone growth is faster in Ezh2^flox/flox^/LysM-Cre^+^ CKO mice

We performed and analyzed micro-CT on 30 days old (right after weaning) tibia trabecular bone of all those genotypic male and female mice. As shown in representative tiabia scans of 30 –day old male mice in Supplemental Fig 1A, among parameters which were analyzed on trabecular bone, we found BV/TV, BS/TV and Tb.N were significantly higher in CKO mice compared with f/+ Cre^-^ and Wt control mice in Supplemental Fig 1B. BV/TV showed significantly higher in f/+ Cre^+^ mice compared with f/+ Cre^-^ and Wt control mice in Supplemental Fig 1B. Tb.Sp showed significantly lower in CKO mice compared with f/+ Cre^+^ and Wt control mice in Supplemental Fig 1B. We did not observe differences of any other parameters in tibia trabecular (Supplemental Table 5), and tibia cortical parameters (Supplemental Table 6). In females, none of those micro-CT parameters were different from the tibia trabecular compartment among groups in Supplemental Fig 1C and 1D and Supplemental Table 7, and there were not statistically different to micro-CT tibia cortical parameters among groups in Supplemental Table 8. We also analyzed micro-CT on 56 days old tibia trabecular bone. In males, BV/TV showed significantly higher in CKO mice compared with f/f, Cre+ and Wt control mice (Supplemental Fig 2A and 2B). BS/TV was significantly higher in CKO mice compared with Wt mice (Supplemental Fig 2B). Tb,N was significantly higher in CKO mice compared with Cre+, f/+ Cre+, f/+ Cre- and Wt mice, but Tb.Sp was significantly lower in CKO mice compared with Cre+ and Wt control mice (Supplemental Fig 2B). Other trabecular micro-CT parameters and cortical parameters were presented in Supplemental Table 9 and 10. In females, BV/TV was significantly lower in CKO mice compared with Cre+, f/+ Cre+, f/+ Cre- and Wt control mice (Supplemental Fig 2C and 2D). In this age of female mice, we found more micro-CT parameters in both trabecular (Supplemental Table 11) and cortical (Supplemental Table 12) in CKO mice to be significantly different compared with other genotypic control mouse groups. From 30 days old young to 6 months old young adults, we did not observe differences on body weight gains between CKO mice and two other control groups (f/f and Cre+) in either males or females (Fig 4A and 4B). The growth of tibia length was also tightly matched in those three groups of mice in both males and females (Fig 4A and 4B). However, in male trabecular bone, BV/TV in CKO mice steadily increased from 30-days-old to 6-months-old, but it went down after 56 days- in f/f and Cre+ control mice (Fig 4C). The changes of TV with age were no differences between CKO and two control mouse groups (Fig 4C). BV and Tb.N were found increased from 30 days old to 56 days old in all three groups, they significantly declined from 56 days old to 6 months old, but such significant declines did not happen in CKO mice (Fig 4C). With regards to cortical bone, the change in BV/TV, BV, B.Ar and Cs.Th from 30 days old to 6 months old were similar across all three groups (Fig 4C). In females, BV/TV gradually increased from 30 days old to 6 months old in CKO mice, however, it increased from 30 days old to 56 days old, and significantly declined from 56 days old to 6 months old in f/f and Cre+ two control mouse groups (Fig 4D). While the differences at 6 months old were described above, the pattern of changes of BV, TV and Tb.N from 30 days old to 56 days to 6 months old were not significantly different among three mouse groups (Fig 4D). On cortical side, the changes of BV/TV, BV, B.Ar and Cs.Th from 30 days old to 6 months old were found similar to we observed in male mice with no differences among all three groups (Fig 4D).

**Figure 4.**
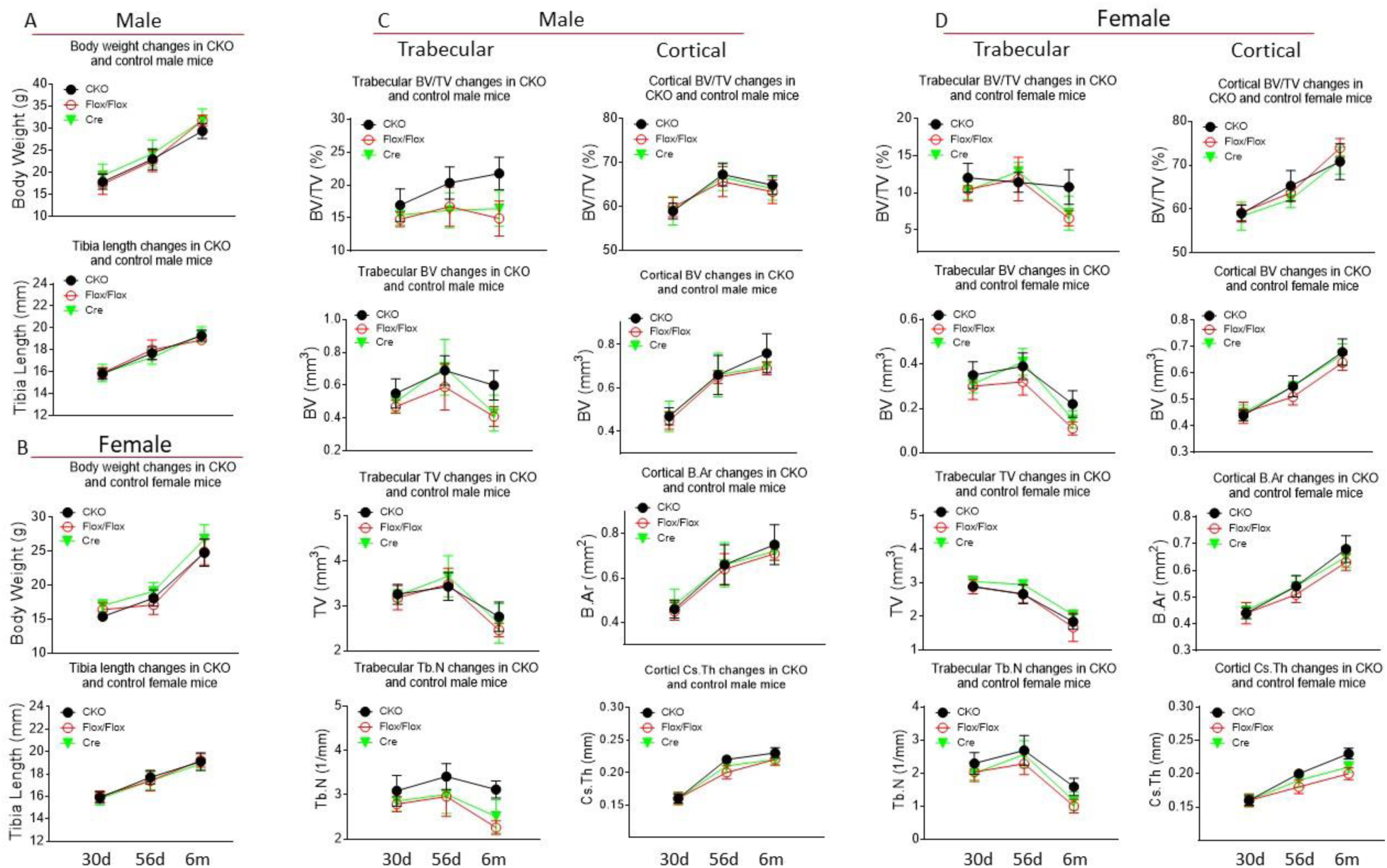
Body weight, tibia length and bone growth from weaning to young adult of male and female Ezh2^flox/flox^/LysM-Cre^+^ CKO and Ezh2^flox/flox^ (Flox/Flox) and LysM-Cre^+^ (Cre) control mice. **(A)** Body weight and tibia length changes from 30 days old to 56 days old to 6 months old of CKO, Flox/Flox and Cre control male mice. **(B)** Body weight and tibia length changes from 30 days old to 56 days old to 6 months old of CKO, Flox/Flox and Cre control female mice. **(C)** Changes of micro-CT trabecular parameter BV/TV, BV, TV and Tb.N, and cortical parameter BV/TV, BV, B Ar (bone area) and Cs Th (cortical thickness) from 30 days old to 56 days old to 6 months old of CKO, Flox/Flox and Cre control male mice. **(D)** Changes of micro-CT trabecular parameter BV/TV, BV, TV and Tb.N, and cortical parameter BV/TV, BV, B Ar (bone area) and Cs Th (cortical thickness) from 30 days old to 56 days old to 6 months old of CKO, Flox/Flox and Cre control female mice. Mouse numbers were used: 30 days old, male: CKO n=9, Flox/Flox n=7, Cre n=9; female: CKO n=8, Flox/Flox n=8, Cre n=8. 56 days old, male: CKO n=9, Flox/Flox n=10, Cre n=10; female: CKO n=10, Flox/Flox n=10, Cre=10. 6 months old, male: CKO n=10, Flox/Flox n=8, Cre n=8; female: CKO n=9, Flox/Flox n=8, Cre n=7.

### Deceased inflammatory milieu and osteoclast numbers in bone of Ezh2^flox/flox^/LysM-Cre^+^ CKO mice at 6 months old

We isolated total proteins from the spine (L4) of CKO, and f/f and Cre+ two control mouse groups at 6 months old, and inflammatory antibody array analysis was performed. In males, almost half of the inflammatory factors measure were significantly lower in CKO mice compared with those from f/f mice (Fig 5A). Among those, M-CSF was significantly lower in CKO mice compared with f/f mice (Fig 5A), notice that M-CSF is a key positive regulator for osteoclast activation and differentiation. When CKO was compared with Cre+ mice, we found more than two third (60%) of those inflammatory factors were significantly lower in CKO mice (Fig 5A). Among those factors, CSFs were also significantly lower in CKO mice compared with Cre+ control mice (Fig 5A). In females, there were 14 out of 40 of those inflammatory factors were significantly lower in CKO mice compared with f/f mice (Fig 5B), and 24 out of 40 (60%) of those inflammatory factors were significantly lower in CKO mice compared with Cre+ control mice (Figure 5B). Notably, in addition to CSF TNFα was also significantly lower in CKO mice compared either f/f or Cre+ control mice (Fig 5B). While the significant differences of expression of some inflammatory factors between CKO and control mice were not profound, these data strongly indicated that inflammatory milieu in bone of both male and female CKO mice was lower compared with their respective control mice, and suggested that octeoclast activity or differentiation is affected by deletion of Ezh2 in osteoclastic cells. We next performed bone histomorphometric analysis to evaluate cellular changes in bone of CKO mice compared with their controls. As shown in Fig 6A, trabecular number and bone area (Fig 6B) were significantly higher in CKO mice compared with either f/f or Cre+ control mice. Bone from CKO mice had less TRAPase positive staining compared with samples from f/f and Cre+ control mice (Fig 6A). Osteoclast number (Fig 6C) and osteoclast surface (Fig 6D) were significantly lower in CKO mice compared with f/f and Cre+ control mice, consistent with findings of the inflammatory array data. Unexpectedly, osteoblast number in bone of Ezh2 osteoclastic cell specific knockout CKO mice significantly higher than it in bone from f/f and Cre+ mice (Fig 6E), suggesting deletion of Ezh2 in osteoclastic cells indirectly affecting on osteoblastogenesis. Consistent to the results we found in males, data from female mice were presented in Supplemental Fig 3. Indeed, when we used Ezh2 and IRF8 antibody double staining on proximal tibia section from CKO and Cre+ mice, we found inverse association between Ezh2 and IRF8 expression on the bone surface of osteoclastic cells (Fig 6F).

**Figure 5.**
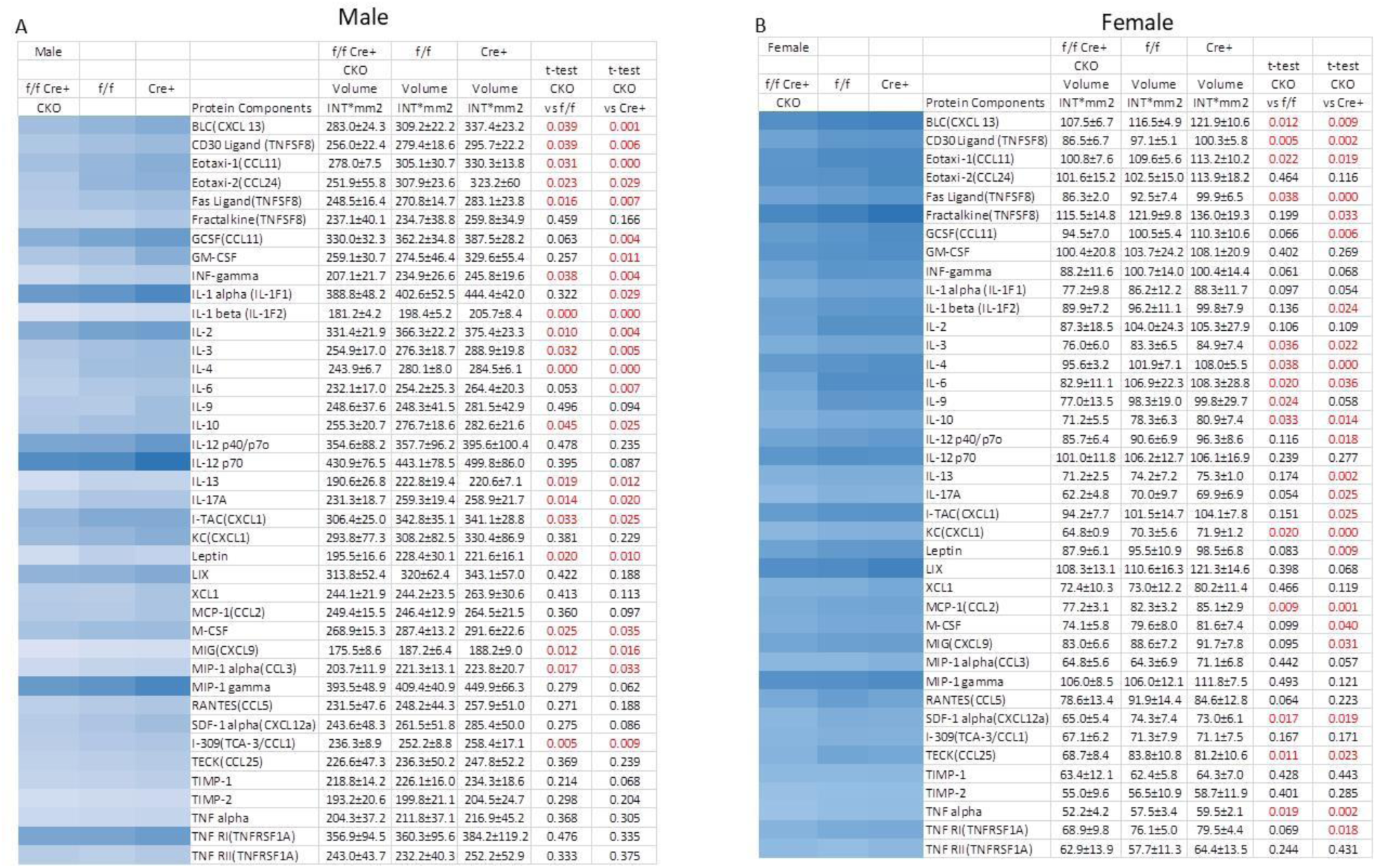
Decreased inflammation in bone from 6 months old male and female Ezh2^flox/flox^/LysM-Cre^+^ (f/f Cre^+^, CKO) compared with Ezh2^flox/flox^ (f/f) and LysM-Cre^+^ (Cre^+^) control mice. **(A)** Inflammatory antibody array analysis showing decreased inflammatory factor expression in male f/f CKO of 6 months old mouse group in total proteins isolated from spine L3 vertebrae compared with those from 6-months-old male f/f and Cre^+^ control mouse groups, the heat map analysis on left side for comparison of all factors. Data are expressed as mean ± SD of blot intensity with samples from each group were pooled to 3 per group, ttest was performed with red showing significance at p<0.05. **(B)** Inflammatory antibody array analysis showing decreased inflammatory factor expression in female f/f CKO of 6 months old mouse group in total proteins isolated from spine L3 vertebrae compared with those from 6-months-old female f/f and Cre^+^ control mouse groups, the heat map analysis on left side for comparison of all factors. Data are expressed as mean ± SD of blot intensity with samples from each group were pooled to 3 per group, ttest was performed with red showing significance at p<0.05.

**Figure 6.**
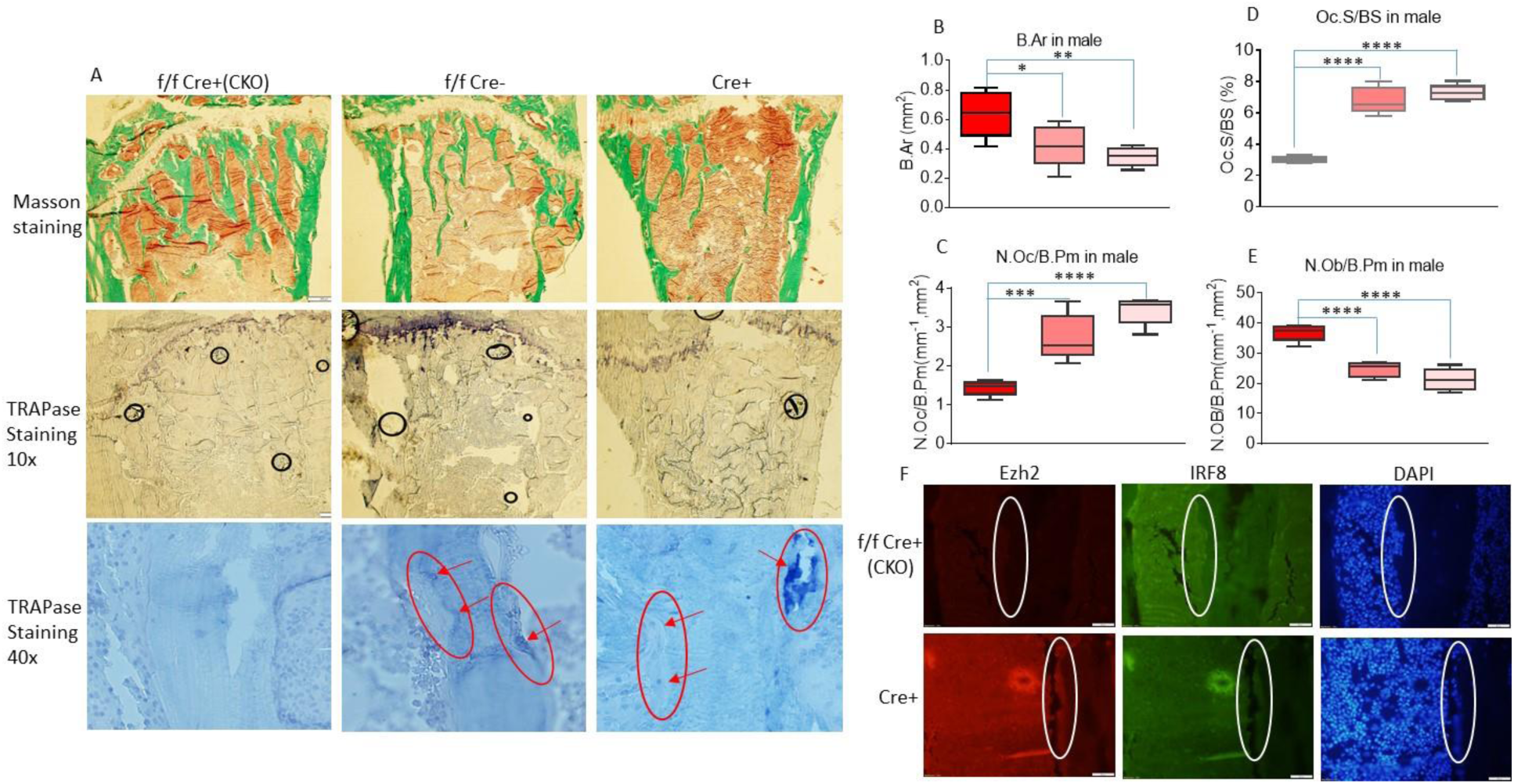
Histomorphometric analysis of tibia of 6 months old male Ezh2^flox/flox^/LysM-Cre^+^ (f/f Cre^+^, CKO) compared with Ezh2^flox/flox^ (f/f Cre^-^) and LysM-Cre^+^ (Cre^+^) control mice. **(A)** Top panel showing Masson staining of proximal tibia below growth plate of representative pictures from each group, green color indicating trabecular and cortical bone; middle panel showing TRAPase staining of proximal tibia below growth plate of representative pictures from each group, dark purple color indicating TRAPase positively stained cells; bottom panel showing TRAPase staining in highest magnification (40x) of one typical osteoclast active area (red circled and arrow pointed) of bone from each group. **(B), (C), (D) and (E)** Parameters of histomorphometric analysis of tibia from f/f Cre^+^ (CKO), f/f and Cre^+^ control mice. B Ar, bone area, N Oc/B Pm, number of osteoclast per bone parameter, Oc S/BS, osteoclast surface per bone surface, N Ob/B Pm, number of osteoblast per bone parameter. Red box bar, CKO group; pink box bar, f/f control group; light pink box bar, Cre control group. Data are expressed as mean ± SD, analyzed by one-way ANOVA, additionally, * p<0.05, ** p<0.01, *** p<0.001, **** p<0.0001 by Tukey’s multiple comparison. Real-time PCR shows β-Catenin and Runx2 mRNA expression in bone from male and female separated samples. **(F)** Ezh2 and IRF8 antibody immune-counterstaining of a representative sample of tibia from either CKO or Cre mouse, 40x on one osteoclast active area white circled.

### Ezh2 controls osteoclast suppressive gene expression regulating osteoclastogenesis

Western blots were performed using total protein isolated from L4 of CKO and f/f and Cre+ mice at 6-monthof age. Two osteoclast bone resorptive markers Cathepsin K and MMP9 weresignificantly lower of their protein expression in CKO mice compared with f/f or Cre+ control mice (Fig 7A). On the other hand, protein levels of osteoclast suppression markers including IRF8, MafB and Arg1 were significantly higher in bone from CKO mice compared with f/f or Cre+ mice (Fig 7A). Unexpectedly, the protein levels of the osteoblastic bone formation markers osteocalcin and Col1 were significantly higher in bone from CKO mice compared with f/f and Cre+ mice (Fig 7A). Serum bone remodeling marker measurements were performed using ELISA. Serum CTX1 and TRAP-5b levels were significantly lower in CKO mice compared with f/f and Cre+ mice (Fig 7B). On the other hand, serum P1NP and osteocalcin levels were significantly higher in CKO mice compared with f/f and Cre+ control mice (Fig 7B). These Western blot and serum bone remodeling marker measurement results found in males were consistent with results found in female mice (Supplemental Fig 4). Indeed, when we isolated total RNA from bone, real-time PCR showed that NFATc1 and Cathepsin K mRNA expression were significantly lower in CKO mice compared with f/f and Cre+ mice (Fig 7C). On the other hand, IRF8, MafB and Arg1 those osteoclast inhibitory factor mRNA and osteocalcin mRNA expression were significantly higher in CKO mice compared with f/f and Cre+ mice (Fig 7C). Moreover, using genomic DNA and chromatins isolated andprecipitated from bone of CKO, f/f and Cre+ mice, we performed H3K27me3 and Ezh2 ChIP analysis. We found that IRF8, MafB and Arg1 those osteoclast inhibitory factors were significantly associated with H3K27me3 in CKO mice compared with f/f and Cre+ mice, but less associated with Ezh2 (Fig 7D). Notice that CKO mice were Ezh2 gene deleted only in osteoclastic cell population, chromatins were precipitated from multi mixed cell types.

**Figure 7.**
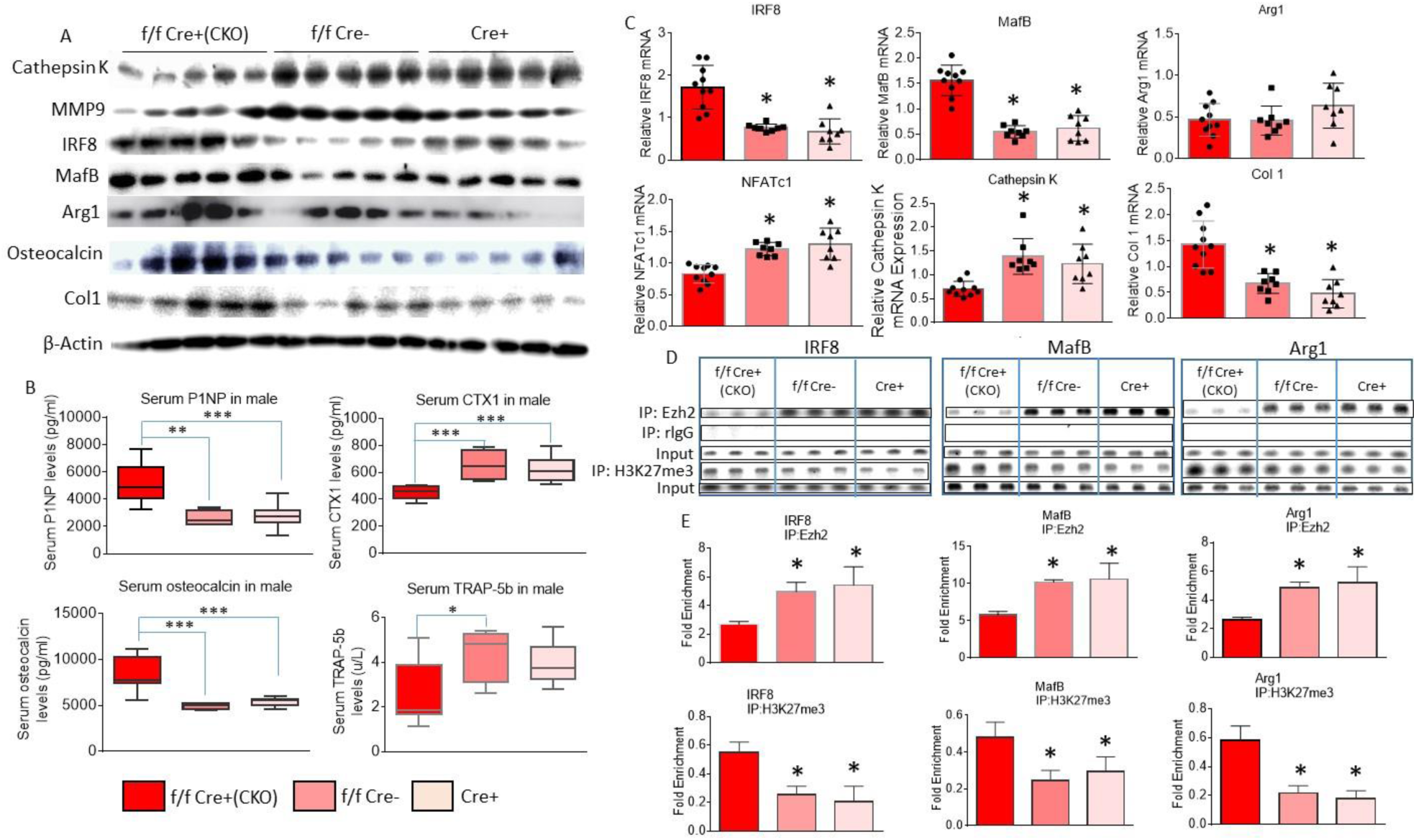
Increased osteoclast suppressive gene expression and decreased bone resorption in 6 months old male Ezh2^flox/flox^/LysM-Cre^+^ (f/f Cre^+^, CKO) compared with Ezh2^flox/flox^ (f/f Cre^-^) and LysM-Cre^+^ (Cre^+^) control mice. **(A)** Western blots showing cathepsin K, MMP9, IRF8, MafB, Arg1, osteoclalcin and Col1 (collagen 1a) expression in total protein isolated from L3 of f/f Cre^+^ CKO, and f/f Cre^-^ and Cre^+^ control mice, samples from each group were pooled to 5 per group. **(B)** Serum ELISA for bone remodeling markers P1NP, osteocalcin, CTX1 and TRAP-5b of f/f Cre^+^ CKO, and f/f Cre^-^ and Cre^+^ control mice. **(C)** Real-time PCR for mRNA expression of IRF8, MafB, Agr1, NFATc1, Cathepsin K and Col1 in bone of total RNA isolated from L3 of f/f Cre^+^ CKO, and f/f Cre^-^ and Cre^+^ control mice. Data are expressed as mean ± SD, analyzed by one-way ANOVA, additionally, * p<0.05, ** p<0.01, *** p<0.001 by Tukey’s multiple comparison. **(D)** Standard ChIP analysis, chromatin from bone was immune-precipitated (IP) by Ezh2 and H3K27me3 and then their associations with IRF8, MafB and Agr1. Samples were pooled to 3 per group. **(E)** Fold changes of IRF8, MafB and Agr1 associated with Ezh2 and H3K27me3 precipitations from standard ChIP assay.

We used pharmacologic inhibitors of Ezh2 to investigate osteoclastogenesis and inverse association of Ezh2 and IRF8 expression using *in ex vivo* bone marrow non-adherent hematopoietic cell and RAW246.7 cell cultures. We used commercially available Ezh2 inhibitors GSK126, GSK343 and DZNep which inhibited osteoclastogenesis *in ex vivo* non-adherent bone marrow hematopoietic cell cultures in a dose dependent manner (Fig 8A and 8B). GSK343 at a concentration of 0.5 µM did inhibit Ezh2 expression (Fig 8C and 8D), when we looked at a single osteoclastic cell which Ezh2 was inhibited (Figure 8C), but IRF8 mRNA expression was significantly higher in RAW246.7 cell cultures (Fig 8D). These data once again indicated that Ezh2 and osteoclast inhibitory factors has inverse relationships of their expression and osteoclast inhibitory factor expression is under epigenetic control through Ezh2 to regulate osteoclastogenesis and bone resorption.

**Figure 8.**
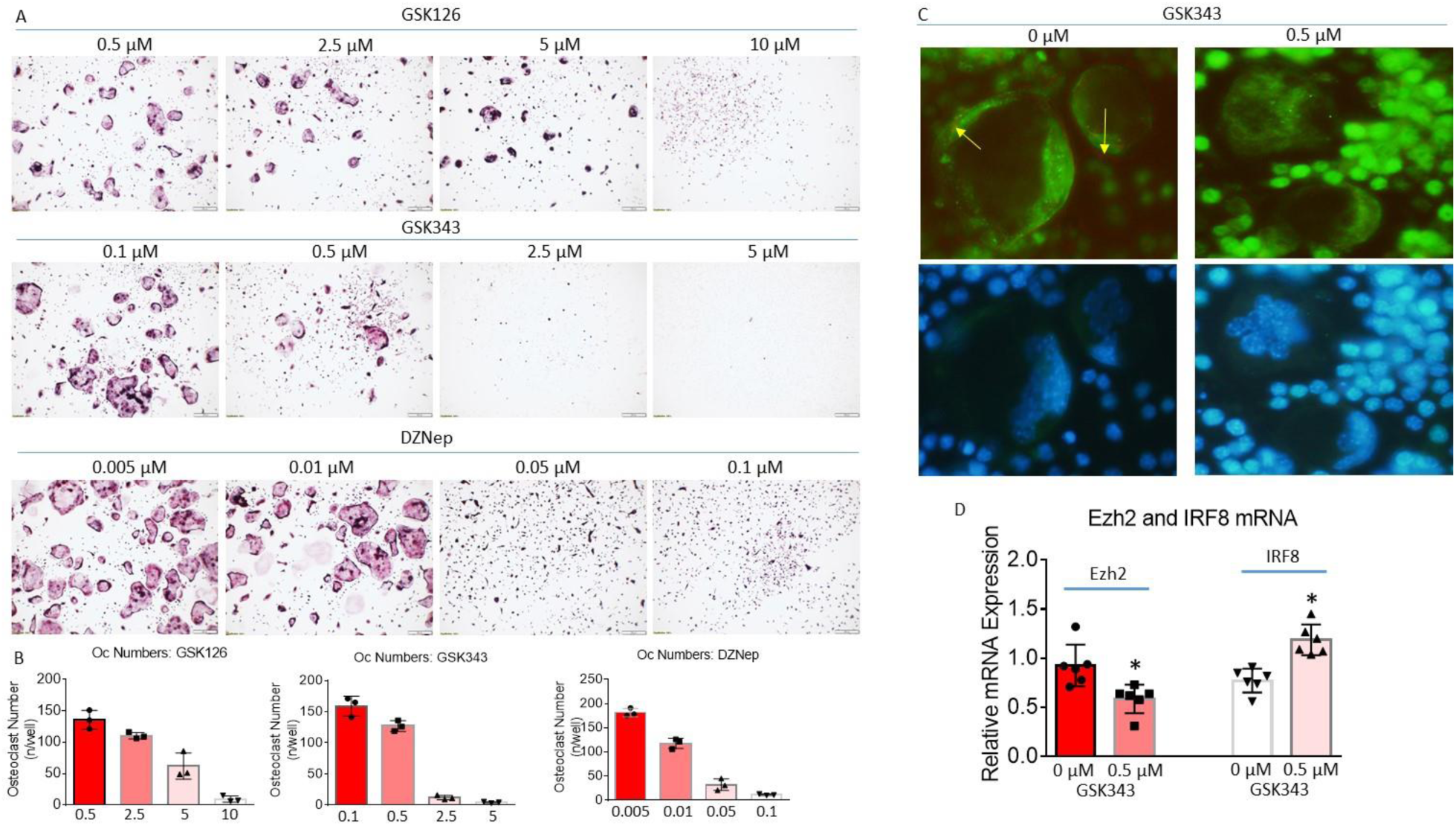
Ezh2 inhibitors suppress osteoclastogenesis. (A) Bone marrow non-adherent hematopoietic cells were isolated from 4-weeks-old wild type male mice. Cells were treated with three different Ezh2 inhibitors GSK126, GSK343 and DZNep with four different concentrations in the presence of 30 ng/ml of RANKL for 5 days. Representative pictures are TRAPase stringing of osteoclast morphology of each Ezh2 inhibitor of each concentration triplicates. (B) Multi-nuclear osteoclast numbers were counted. (C) RAW246.7 cells were treated with or without 0.5 µM of GSK343 in the presence of 30 ng/ml of RANKL for 5 days, cells were then immune counter-stained with Ezh2 (red) and IRF8 (green) antibodies. Upper pane yellow arrow showing Ezh2 red stained osteoclast, low panel is DAPI staining. (D) RAW246.7 cells were treated with or without 0.5 µM of GSK343 in the presence of 30 ng/ml of RANKL for 5 days. RNA was isolated for real-time PCR of Ezh2 and IRF8 mRNA expression. * p<0.05 ttest.

## Discussion

In our current report, we have determined that deletion of Ezh2 gene in osteoclast precursors increases bone mass in both male and female mice. Densitometric and structural analysis on cortical compartment showed bone was not osteopetrotic in Ezh2 CKO mice, indicating that Ezh2 is not essential for late stage of osteoclast differentiation, but it significantly contributes to controlling osteoclstogenesis and bone resorption in young adult mice. Postnatal bone accretion was accelerated in mice lacking Ezh2. In addition, Ezh2 deletion differential influences bone mass acquisition and bone homeostasis in weanling and adult mice. Moreover, we provide evidence of sxual dimorphism with regards to Ezh2 function in bone development and turnover. In males, not only CKO mice but also Ezh2^flox/+^/LysM-Cre+ genotypic mice showed increased bone mass, while in females only CKO mice had increased bone mass phenotype compared with other genotypic young adult control mice.

Our study provide new mechanistic data concerning the role of Ezh2 in osteoclast function and bone mass accretion. Specifically, in the absence or inactivated Ezh2 in pre-osteoclasts, osteoclastic suppressive genes are activated leading to suppression of osteoclastogenesis and osteoclast activity. This suggests that osteoclastogenesis and bone resorption are under epigenetic regulation through Ezh2. On the other hand, deletion of Ezh2 in osteoclast precursors also inhibited genes of those known as osteoclastogenic or required for osteoclast differentiation and activity, but indirectly activated osteoblast markers. These data suggest that Ezh2 gene expression plays a key role in governing osteoclast signaling, thereby altering the balance between bone resorption and bone formation. An altered profile of inflammatory markers in pre-osteoclastic cells from mice lacking Ezh2 may provide an environmental signal for osteoclast and osteoblast differentiation and activity. Collectively, our data identify Ezh2 as a potential target for strategies aimed at treating both osteoclastic bone resorptive disorders such as osteoporosis and rheumatoid arthritis.

Modification of osteoclast gene expression through histone modification during osteoclastogenesis might be a good approach to control osteoclast differentiation or activity and thus alter osteoclastic bone resorption (Yi *et al*, 2019). Osteoclasts are large multinucleated cells with ruffled membranes, and they are differentiated from mononuclear cells of the hematopoietic stem cell lineage by fusing those monocytes to form tartrate-resistant acid phosphatase (TRAP)-positive cells (Levaot *et al*, 2015). It has been known for many years that two factors, macrophage colony-stimulating factor (M-CSF) and receptor activator of nuclear factor kappa-B ligand (RANKL) stimulate osteoclast differentiation, and RANKL is essential for the formation, fusion, activation and survival of osteoclasts (Asagiri & Takayanagi, 2007). Now, there is a growing body of evidence suggested that histone methyltransferase is involved in bone cell differentiation (Cakouros & Gronthos, 2019). Ezh2 has been reported to suppress osteoblastogenesis via H3K27me3 methylation on the promoters of osteoblast-related genes (Wei *et al*, 2011). On the other hand, *in vitro* studies have shown that Ezh2 promotes osteoclastogenesis by downregulating IRF8, a negative regulator of osteoclastogenesis (Adamik *et al*, 2020). Studies have also shown that H3 monomethylation at lysine 27 by G9a is essential for the enzymatic activity of MMP-9 and facilitates osteoclast differentiation (Shin *et al*, 2019), and DOT1L a histone methyltransferase of H3K79me suppressed osteoclast differentiation (Gao & Ge, 2018). In our current report, we have successfully generated a pre-osteoclast, macrophage specific Ezh2 gene deletion mouse model. Before mice reach to 56 days old, bone development of those CKO mice seem to be no significantly different compared with their control littermates. Compared with all other genotypic littermates generated by in-bred breeding, Ezh2 deletion in pre-osteoclastic LysM-Cre positive cells promotes bone mass accretion. Indeed, we provide clear evidence *in vivo* that deletion of Ezh2 in pre-osteoclastic cells inhibits osteoclastogenesis in young adult mice (Chen *et al*, 2020).

Ezh2 is known as a histone lysine methyltransferase associated with predominantly transcriptional or gene repression (Nutt *et al*, 2020), it catalyzes the addition of methyl groups to H3K27me3 in target genes involved in numerous cellular processes. We have originally showed evidence that Ezh2 might be involved in bone cell differentiation in the context of maternal or early life diet-induced obesity (Chen *et al*, 2016). Although our current studies focused on determining the role of Ezh2 in osteoclastogenesis and bone resorption, we have previously investigated the role of Ezh2 in the function of osteoblastic cells. These data suggest that maternal HFD-induced obesity increases Ezh2 expression in fetal osteoprogenitors, while increased Ezh2 activity suppressed expression of osteoblastic marker genes such as SATB2. Ezh2 facilities association of H3K27me3 with SATB2 in fetal cells from both obese dams in rodents and obese mothers in human (Chen *et al*, 2020). We described that deletion of Ezh2 gene in osteoblastic specific cell lineage increased bone density, especially trabecular bone density and bone mineral content, and increased SATB2 expression in bone (Chen *et al*, 2020). Together with data presented in our current report, these results clearly demonstrats that Ezh2 expression controls both osteoblast and osteoclast differentiation and activity. Repressive H3K27me3 and active H3K27ac are epigenetic marks that control gene expressions associated with them, some of them are essential in controlling osteoclast and osteoblast differentiation (Yi *et al*, 2019), however, there are only few studies offer little and inconsistent insight on how epigenetic remodeling of bone-specific chromatin maintains bone cell development and bone mass *in vivo*. Genetic inactivation of Ezh2 in calvarial bones using Ezh2^flox/flox^ and Prrx1-Cre model enhanced expression of osteogenic genes (Dudakovic *et al*, 2020). Evidence from several other studies also support a role for Ezh2inhibition in stimulating bone accretion (Ferguson *et al*, 2018). It has been shown that loss or inhibition of Ezh2 not only resulted in enhanced osteogenesis, but also inhibited adipogenic differentiation of mesenchymal stem cells from both human and rodents (Hemming *et al*, 2014; Lui *et al*, 2016). These were interesting evidence describing the role of Ezh2 on bone cell development, however, bone development and modeling are controlled by both osteoblastic and osteoclastic cells, therefore, to determine how Ezh2 controls osteoclast development *in vivo* is important. However, we have only seen few evidence from *in vitro* or *in ex vivo* studies demonstrated the role of Ezh2 during osteoclastogenesis (Adamik *et al*, 2020). We have shown that deletion of Ezh2 in pre-osteoclastic cells resulted in reduced trimethylation in those genes that play suppressive functions on osteoclast differentiation. Interestingly, for those genes that are responsible for stimulating osteoclast were down-regulated.

Osteoclastic suppressive genes have been identified such as IRF8 (interferon regulatory factor 8), MafB (V-mad musculoaponeurotic fibrosarcoma oncogene homologue B) and Arg1 (Arginase 1), IRF8 has received significant attention lately due to its role in inhibiting osteoclast differentiation (Thumbigere-Math *et al*, 2019). It has recently been reported that both homo and heterozygous of IRF8 gene mutations promote osteoclastogenesis and increased osteoclast activity, which is accompanied by increased NFATc1 expression (Izawa *et al*, 2019). IRF8 plays critical roles in the regulation of lineage commitment and in myeloid cell maturation including the decision for a common myeloid progenitor to differentiate into a monocyte precursor cell (Das *et al*, 2020). The LysM-Cre mouse model used in our current study has a nuclear-localized Cre recombinase inserted into the first coding ATG of the lysozyme 2 gene (Lyz2); both abolishing endogenous Lyz2 gene function and placing NLS-Cre expression under the control of the endogenous Lyz2 promoter/enhancer elements (The Jackson Laboratory). These LysM-Cre mice were generated to be used for Cre-lox studies of the myeloid cell lineage (monocytes, mature macrophages and granulocytes) and the innate immune response. Therefore, we expected that Ezh2 gene deletion in the myeloid cell lineage could interfere with IRF8 expression and function. We have showed that Ezh2 gene deletion resulted in increased IRF8 gene expression, therefore osteoclastogenesis and bone resorption were inhibited. Indeed, ChIP assay indicated that Ezh2 and IRF8 are highly associated, and Ezh2 controls IRF8 gene trimethylation are obviously through H3k27me3. Further, overexpression of Ezh2 in osteoclastic cells (murine macrophage cell line RAW264.7) suppresses IRF8 expression, and osteoclastogenesis was increased. These data suggest that Ezh2 and IRF8 axis in the myeloid cell lineage is at least one of mechanisms of how Ezh2 controls osteoclastogenesis and bone resorption.

Deletion of Ezh2 in pre-osteoclasts resulted in a significant reduction in activities of inflammatory factors within bone. We do not have explanation for why Ezh2 deletion alters the bone inflammasome, we only speculate that inflammatory signatures in macrophages in bone can be programmed through Ezh2 expression. Although decreased osteoclastogenesis and increased bone mass in 6 months old Ezh2 lacking male and female mice are clear, the postnatal bone modeling and growth in young CKO mice were more ambiguous. However, we did not observe obvious differences of bone mass in 30 days old CKO mice compared with all other genotypic and wild type mice in both males and females. Since bone mass in young adult CKO mice was significantly greater than control mice, overall bone growth in CKO mice was faster from 30 days old to 6 months old. The reason for why 30 days old CKO mice showed no obvious bone phenotypic changes compared to control may be because, 1) Ezh2 deletion is in pre-osteoclastic cells, 2) 30 days old mice were just after weaning, osteoblastic bone formation is a dominate event for rapid bone growth during this age of mice (Chen *et al*, 2017). Moreover, the function of osteoclasts during rapidly growing phase are probably to make the bone in a good shape (Chen *et al*, 2017), but not much on bone mass. This is supported by our data, although overall there were no statistically different among groups on bone density, but trabecular centroid (x) in 30 days old CKO mice was significantly different compared to their control littermates (Cre+ mice) in both sexes. We stated that CKO mice were notosteopetrotic. Osteopetrosis is defined by increased bone density due to a defect in bone resorption by osteoclasts. In mice laking Ezh2, we did not observe clear differences in the cortical bone compartment by micro-CT analysis, and there were still osteoclasts in bone of CKO mice. Therefore, our determination is that in CKO mice, osteoclastic bone resorption was decreased but not completely abolished. Lastly, we have shown that in males, not only CKO mice but also Ezh2^flox/+^/LysM-Cre+ genotypic mice showed increased bone mass, while in females only CKO mice had increased bone mass phenotype compared with control mice (compared with either flox, Cre and wild type control) in young adult mice. This observation suggests a potential role for sex hormones or Ezh2 interfered with X chromosome in bone accretion and modeling, which warrants further investigation.

In summary, we show that Ezh2 deletion in osteoclast precursors increased postnatal bone development as well as bone mass in young adult mice. Our data demonstrate a role for Ezh2 in osteoclast gene expression, modulating the bone inflammasome, as well as controlling bone resorption. We suggest that manipulation of Ezh2 expression may be a viable strategy to alter bone development or manage bone resorptive disorders such as osteoporosis and rheumatoid arthritis.

## Materials and Methods

### Animals and diets

C57BL/6J mouse background myeloid cell-specific targeted mutant LysMcre (LysM-Cre^+^) mice (Stock No: 004781) and loxP sites flanking exons 14-15 of the zeste homolog 2 (Ezh2) gene Ezh2^flox/flox^ mice (stock no. 022616) were purchased from the Jackson Laboratory (Bar Harbor, ME). To delete targeted gene of Ezh2 in the myeloid cell lineage of osteoclast precursors, including monocytes, mature macrophages, and granulocytes, LysM-Cre^+^ mice were crossed with Ezh2^flox/flox^ mice. Male Ezh2^flox/+^/LysM-Cre^+^ and female Ezh2^flox/+^/LysM-Cre^-^ offspring were intercrossed to generate the LysM-Cre^+^ (Cre-positive control); Ezh2^flox/flox^ /LysM-Cre^+^ mice (CKO) and others Ezh2^flox/+^/LysM-Cre^-^, Ezh2^flox/flox^/LysM-Cre^-^ (flox control), Ezh2^flox/+^/LysM-Cre^+^, and wild type (Wt) mice. A staggering mouse breeding strategy was used and all mice were fed standard rodent chow diet. Experiments involved 30 days old, 56 days old and 6 months old littermates of male and female CKO and their corresponding genotypic and wild type control male and female mice (8-12 per group) for bone phenotyping studies. Mice were weighed, randomized by their weights, and housed 5 per cage in mouse small shoe box cages. All animal procedures were approved by the Institutional Animal Care and Use Committee at University of Arkansas for Medical Sciences (AUP#3595 UAMS, Little Rock, AR), and housed in animal facility at the Arkansas Children’s Research Institute with constant humidity and lights on from 06:00-18:00 hrs at 22°C. At the end of the studies, mice were anesthetized by injection with 100 mg Nembutal/kg body weight (Avent Laboratories). Blood was collected via cardiac puncture, which was followed by decapitation, femur, tibia and vertebrae bones were collected and stored in -80 °C.

### Bone analyses

Micro-computed tomography (micro-CT) measurements of the trabecular and cortical compartments from the left tibial and L5 spine bone were evaluated using SkyScan μCT scanner (recently upgraded SkyScan 1272, Bruker.com) at 8 μm pixel size with X-ray source power of 60 kV and 166 μA and integration time of 950 ms. The trabecular compartment included a 0.9 mm region extending distally 0.03 mm from the physis. The grayscale images were processed by using a filter (=Al, σ=0.5, mm) to remove noise, and a fixed threshold of 125 was used to extract the mineralized bone from the soft tissue and marrow phase. Cancellous bone was separated from the cortical regions by semi-automatically drawn contours. A total of 120 slices starting from about 0.1 mm distal to growth plate, constituting 0.80 mm length, were evaluated for trabecular bone structure, bone volume fraction (BV/TV, %), trabecular thickness (Tb.Th, mm), trabecular separation (Tb.Sp, mm), trabecular number (Tb.N, 1/mm), Degree of anisotropy (DA) were calculated based on description by Bouxsein et al. (Bouxsein *et al*, 2010), and by using software provided by SkyScan, Bruker. For cortical bone, the cortical compartment was a 0.6 mm region extending distally starting 5 mm proximal to the tibia-fibula junction. Total cross-sectional area (CSA, mm^2^), medullary area (MA, mm^2^) and cortical thickness (Ct.Th, mm) were assessed.

### Bone histology

Mouse right tibia samples were embedded, cut and TRAPase stained by Histology Special Procedures at the Arkansas Children’s Nutrition Center Histology Core. TRAPase staining kit was utilized according to the manufacturer’s protocols (Sigma-Aldrich, Acid phosphatase leukocyte, procedure No. 386). TRAPase positive pink-purple-stained cells were visualized with a digitizing morphometry system, which consists of an epifluorescent microscope (model BH-2, Olympus), a color video camera, and a digitizing pad (Numonics 2206) coupled to a computer (Sony) (OsteoMetrics, Inc.).

### Measurements of bone turnover markers in bone marrow plasma and in serum

Bone marrow plasma were prepared at the time of tissue harvest. Bone marrow was flushed out from femur using 300 µl of PBS, vortexed and spun (1700 g) for collecting supernatant as bone marrow plasma. Serum bone resorption marker C-terminal telopeptides of type I collagen (CTX-1) was measured by Rat-Laps^TM^ ELISA from Nordic Biosciences Diagnostic (Herlev, Denmark). The serum P1NP (Procollagen 1 N-terminal Propeptide) levels were measured by direct immunoassay P1NP assay Kit. The P1NP level measurement ELISA kit was purchased from Mybiosource.com (Catalog No: MBS2500076) and measurement procedure followed the manufacturer’s recommendations. According to the protocol, pre-coated with total-P1NP antibody, total-P1NP present in the sample is added and binds to antibodies coated on the wells, and then biotinylated total-P1NP antibody is added and binds to total-P1NP in the sample. Then, streptavidin-HRP is added and binds to the biotinylated total-P1NP antibody. After incubation unbound streptavidin-HRP is washed away during a washing step, substrate solution is then added and color develops in proportion to the amount of total-P1NP. ELISA for serum osteocalcin and TRAP-5b were performed according to procedure from manufactory. The reaction is terminated by addition of acidic stop solution and absorbance is measured at 450 nm.

### Cell cultures

Murine osteoclast cell line RAW264.7 cells (from ACCT) and non-adherent bone marrow cells were cultured in 96-well plates (2×10^4^ cells/well) in the presence or absence of 30 ng/ml of RANKL, in α-MEM supplemented with 10% fetal bovine serum (FBS) (Hyclone, Logan, UT), penicillin (100 Units/ml), streptomycin (100 µg/ml), and glutamine (4 mM). These cell cultures were previously described in our laboratory (Chen *et al*, 2020). After 5 days for bone marrow cell cultures, the cells were fixed with 4% paraformaldehyde and stained for TRAPase activity using a TRAPase staining kit according to the manufacturer’s protocols (Sigma-Aldrich, Acid phosphatase leukocyte, procedure No. 386). TRAP-positive cells containing >3 nuclei in each well were counted as osteoclasts under an epifluorescent microscope (model BH-2, Olympus, Imaging America Inc.; Center Valley, PA). For RNA and protein expression study, non-adherent bone marrow cells were seeded in triplicate in 6-well collagen-coated plates (BD Biosciences) at a density of 1×10^5^ cells/well, and cells were treated with Ezh2 inhibitors GSK126, GSK343 and DZNep at four different concentrations in the presence of 30 ng/ml RANKL for 5 days. Murine bone marrow mesenchymal stromal cell line ST2 cells were culture in the osteoblast differentiation medium.

### RNA isolation, real-time reverse transcription-polymerase chain reaction

Bone and cell RNA isolation was performed using TRI Reagent (MRC Inc., Cincinnati, OH) according to the manufacturer’s recommendations followed by DNase digestion and column cleanup using QIAGEN mini columns (Chen *et al*, 2018). Reverse transcription was carried out using an iScript cDNA synthesis kit from Bio-Rad (Hercules, CA). All primers for real-time PCR analysis used in this report were designed using Primer Express software 2.0.0 (Applied Biosystems).

### Western blotting

Total protein extracts from L4 and culture cells were prepared using radioactive immunoprecipitation assay (RIPA) buffer (Solarbio). Western blots were performed using standard protocols (Chen *et al*, 2018). The protein lysates were quantified and separated by sodium dodecyl sulfate polyacrylamide gel electrophoresis and transferred to polyvinylidene fluoride membranes (Millipore). This was followed by immunoblotting with primary antibodies NFATc1 (SAB2101576, Sigma-Aldrich, St. Louis, MO, USA), MMP9 (sc-6840, Santa Cruz, Dallas, TX, USA), Cathepsin K (sc-48353, Santa Cruz, Dallas, TX, USA), Ezh2 (Cell Signaling, #9346), IRF8 (Invitrogen, #39-8800), Arg1 (Cell Signaling, #93668), MafB (Bethyl, A700-046), Osteocalcin (Millipore, AB10911), Col1 (C2456, Sigma-Aldrich, St. Louis, MO, USA) and β-Actin (A1978, Sigma-Aldrich, St. Louis, MO, USA); 1:1000 dilution and then by the corresponding horseradish peroxidase conjugated secondary antibodies. Blots were developed using chemiluminescence (PIERCE Biotechnology) according to the manufacturer’s recommendations. Bands of interest were visualized and imaged under chemiluminescent detection using a FluorChem E System (ProteinSimple, San Jose, CA). Quantitation of the intensity of the bands in the autoradiograms was performed using a VersaDocTM imaging system (Bio-Rad).

### Chromatin-immunoprecipitation (ChIP)

The procedure for standard ChIP assay using Ezh2 (Cell Signaling, #9346) and H3K27me antibody (ChIP grade from Cell Signaling, #07-449) has been described previously (Chen *et al*, 2018), and information for all primers used for ChIP assay and real-time PCR were designed using Primer Express software 2.0.0 (Applied Biosystems). The procedure for standard ChIP assay using H3K27me3 and Ezh2 (Cell Signaling) antibodies was described previously (Chen *et al*, 2018).

### Inflammatory antibody array

Simultaneous detection of multiple cytokines undoubtedly provides a powerful tool to study cell signaling pathways. RayBio C-Series mouse (#AAM-INF-1-8, RayBiotech, Inc) inflammatory antibody arrays were performed for the semi-quantitative detection of 40 mouse proteins in cell lysate according to protocols provided by the company. L4 vertebra bone tissue proteins were extracted using a cell lysate buffer as described previously (Chen *et al*, 2016). Total protein 50 µg from each sample which the original lysate concentration ranged 1-5 mg/ml was loaded. According to description provided by the company, this array can detect IL-2 at concentration of 25 to 250,000 pg/ml, a range of 10,000-fold, and as determined by densitometry, as little as 4 pg/ml of MCP-1 can be detected, the inter-array Coefficient of Variation (CV) of spot signal intensities is 5-10%. Quantitation of the intensity of the bands in the autoradiograms was performed using a VersaDoc^TM^ imaging system (Bio-Rad).

### Statistics

Numerical variables were expressed as means ± SD (Standard Deviation), and n represents the number of samples/group. For *in vitro* and *ex vivo* experiments, differences within groups were evaluated using t test or one-way ANOVA followed by Tukey’s post hoc test. P<0.05 was considered significant. Cell culture experiments were conducted at least three independent times, and representative images (including bone scan images) are displayed. Dose or time response was assessed using Cruick’s non-parametric test for trend. For *in vivo* animal experiments, comparisons among groups were performed with the nonparametric Kruskall-Wallis test followed by a Dunnett’s test comparing each genotype group to the control or CKO group. The nonparametric Wilcoxon rank-sum test was used to compare CKO to individual LysM-Cre positive or Ezh2 flox and other genotypic and wild type mouse groups. Values were considered statistically significant at p<0.05.

## Author contributions

J.R.C. designed and performed the study, and wrote the paper; O.P.L. and M.L.B. performed cell, biochemical and molecular experiments, and *in vivo* sample analysis, helped to perform ChIP analysis; D.G. and C.L. performed *in ex vivo* and *in vitro* experiments; and F.Z. contributed in the performance of experiments and study designs.

## Acknowledgement

This work was supported by sub-objective to J.R.C. by United States Department of Agriculture (USDA) / Agricultural Research Service (ARS) Project # USDA-ARS Project 6026-51000-012-06S to the Arkansas Children’s Nutrition Center. Authors would like to thank Jim Sikes, Matt Ferguson, Hoy Pittman and Bobby Fay for their technical assistances on animal experiments.

---

**Supplemental Figure 1.**
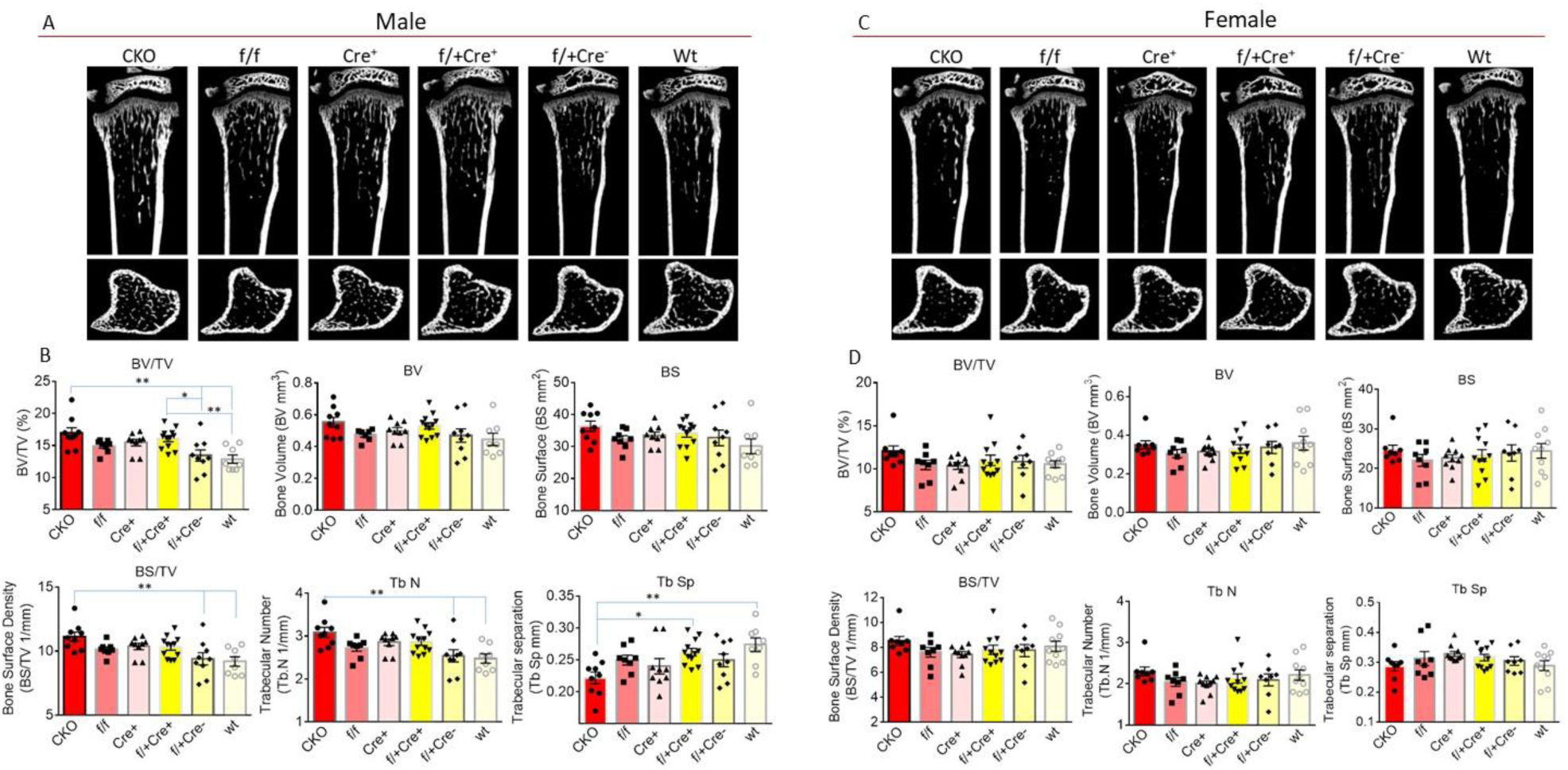
Micro-CT bone phenotype of tibia of 30 days old male and female Ezh2^flox/flox^/LysM-Cre^+^ CKO and other genotypic and wild type control mice. **(A)** Representative micro-CT images of male proximal tibia from one sample from each group of mice, upper panel shows sagittal view and lower panel shows transverse view, white lines and dots indicates trabecular or cortical bone tissues. **(B)** Micro-CT measures of six parameters from trabecular tibias from CKO (Ezh2^flox/flox^/LysM-Cre^+^, n=10), f/f (Ezh2^flox/flox^ control, n=8), Cre^+^ (LysM-Cre^+^ control, n=8), f/+Cre^+^ (Ezh2^flox/+^/LysM-Cre^+^, n=9), f/+Cre^-^ (Ezh2^flox/+^/LysM-Cre^-^, n=7) and Wt (wild type control, n=9) male mice. BV/TV, bone volume/total tissue volume; BV, bone volume; BS, bone surface; BS/TV, bone surface density; Tb.N, trabecular number; Tb.Sp, trabecular separation. **(C)** Representative micro-CT images of female proximal tibia from one sample from each group of mice, upper panel shows sagittal view and lower panel shows transverse view, white lines and dots indicates trabecular or cortical bone tissues. **(D)** Micro-CT measures of six parameters from trabecular tibias from CKO (n=9), f/f (n=8), Cre^+^ (n=7), f/+Cre^+^ (n=8), f/+Cre^-^ (n=8) and Wt (n=8) female mice. Parameter BV/TV, BV, BS, BS/TV, Tb.N and Tb.Sp were presented. Data are expressed as mean ± SD, analyzed by one-way ANOVA, additionally, * p<0.05, ** p<0.01 by Tukey’s multiple comparison.

**Supplemental Figure 2.**
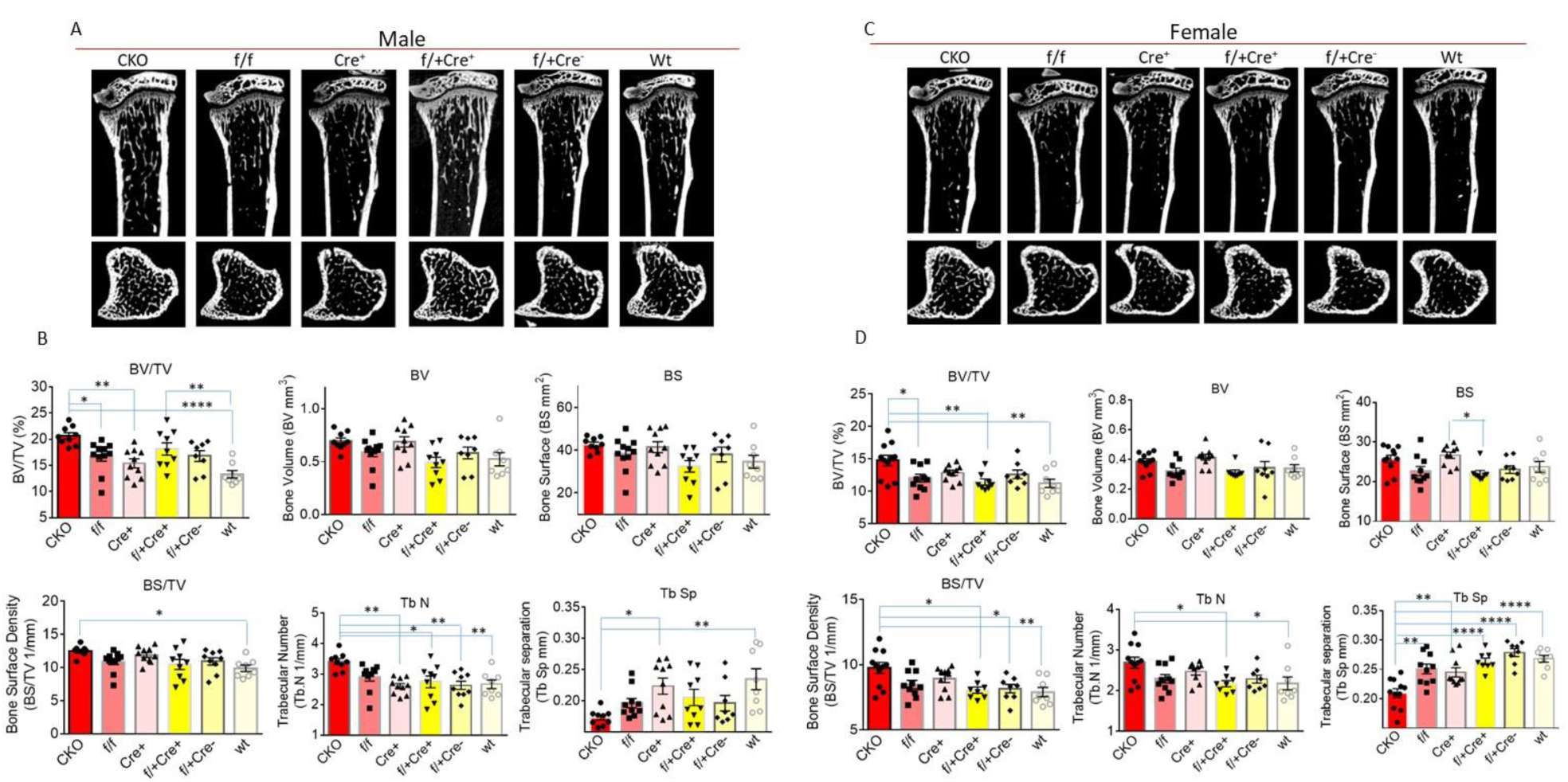
Micro-CT bone phenotype of tibia of 56 days old male and female Ezh2^flox/flox^/LysM-Cre^+^ CKO and other genotypic and wild type control mice. **(A)** Representative micro-CT images of male proximal tibia from one sample from each group of mice, upper panel shows sagittal view and lower panel shows transverse view, white lines and dots indicates trabecular or cortical bone tissues. **(B)** Micro-CT measures of six parameters from trabecular tibias from CKO (Ezh2^flox/flox^/LysM-Cre^+^, n=10), f/f (Ezh2^flox/flox^ control, n=8), Cre^+^ (LysM-Cre^+^ control, n=8), f/+Cre^+^ (Ezh2^flox/+^/LysM-Cre^+^, n=9), f/+Cre^-^ (Ezh2^flox/+^/LysM-Cre^-^, n=7) and Wt (wild type control, n=9) male mice. BV/TV, bone volume/total tissue volume; BV, bone volume; BS, bone surface; BS/TV, bone surface density; Tb.N, trabecular number; Tb.Sp, trabecular separation. **(C)** Representative micro-CT images of female proximal tibia from one sample from each group of mice, upper panel shows sagittal view and lower panel shows transverse view, white lines and dots indicates trabecular or cortical bone tissues. **(D)** Micro-CT measures of six parameters from trabecular tibias from CKO (n=9), f/f (n=8), Cre^+^ (n=7), f/+Cre^+^ (n=8), f/+Cre^-^ (n=8) and Wt (n=8) female mice. Parameter BV/TV, BV, BS, BS/TV, Tb.N and Tb.Sp were presented. Data are expressed as mean ± SD, analyzed by one-way ANOVA, additionally, * p<0.05, ** p<0.01, *** p<0.001, **** p<0.0001 by Tukey’s multiple comparison.

**Supplemental Figure 3.**
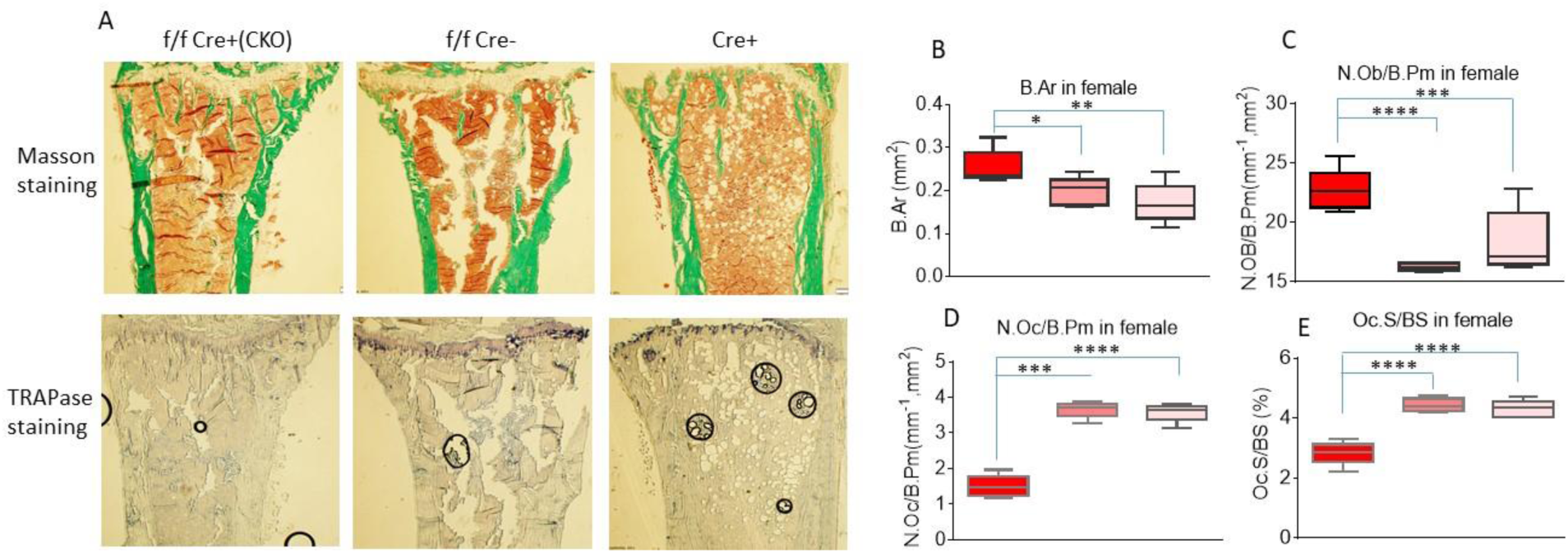
Histomorphometric analysis of tibia of 6 months old female Ezh2^flox/flox^/LysM-Cre^+^ (f/f Cre^+^, CKO) compared with Ezh2^flox/flox^ (f/f Cre^-^) and LysM-Cre^+^ (Cre^+^) control mice. **(A)** Top panel showing Masson staining of proximal tibia below growth plate of representative pictures from each group, green color indicating trabecular and cortical bone; bottom panel showing TRAPase staining of proximal tibia below growth plate of representative pictures from each group (10x), dark purple color indicating TRAPase positively stained cells; **(B), (C), (D) and (E)** Parameters of histomorphometric analysis of tibia from f/f Cre^+^ (CKO), f/f and Cre^+^ control mice. B Ar, bone area, N Ob/B Pm, number of osteoblast per bone parameter, N Oc/B Pm, number of osteoclast per bone parameter, Oc S/BS, osteoclast surface per bone surface, Red box bar, CKO group; pink box bar, f/f control group; light pink box bar, Cre control group. Data are expressed as mean ± SD, analyzed by one-way ANOVA, additionally, * p<0.05, ** p<0.01, *** p<0.001, **** p<0.0001 by Tukey’s multiple comparison.

**Supplemental Figure 4.**
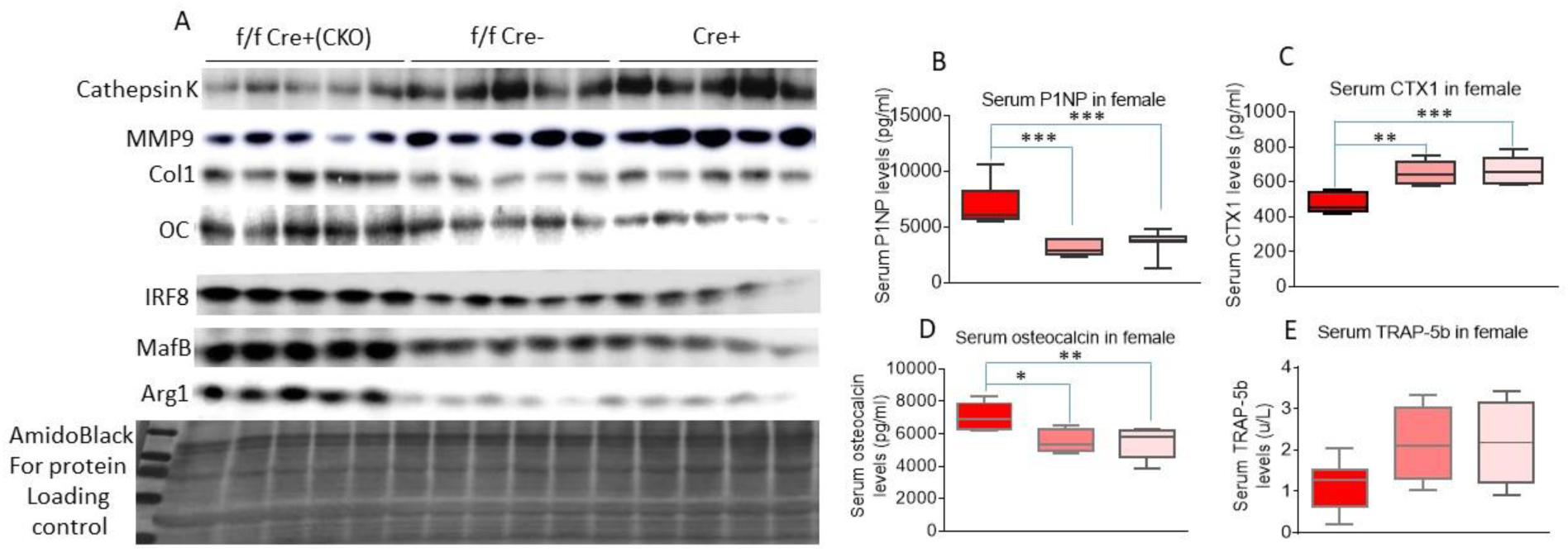
Increased osteoclast suppressive protein expression and decreased bone resorption in 6 months old female Ezh2^flox/flox^/LysM-Cre^+^ (f/f Cre^+^, CKO) compared with Ezh2^flox/flox^ (f/f Cre^-^) and LysM-Cre^+^ (Cre^+^) control mice. **(A)** Western blots showing cathepsin K, MMP9, Col1 (collagen 1a), osteoclalcin (OC), IRF8, MafB, Arg1 expression in total protein isolated from L3 of f/f Cre^+^ CKO, and f/f Cre^-^ and Cre^+^ control mice, samples from each group were pooled to 5 per group. **(B) (C) (D) (E)** Serum ELISA for bone remodeling markers P1NP, CTX, osteocalcin and TRAP-5b of f/f Cre^+^ CKO, and f/f Cre^-^ and Cre^+^ control mice. Data are expressed as mean ± SD, analyzed by one-way ANOVA, additionally, * p<0.05, ** p<0.01, *** p<0.001 by Tukey’s multiple comparison.

**Supplemental Table 1.**
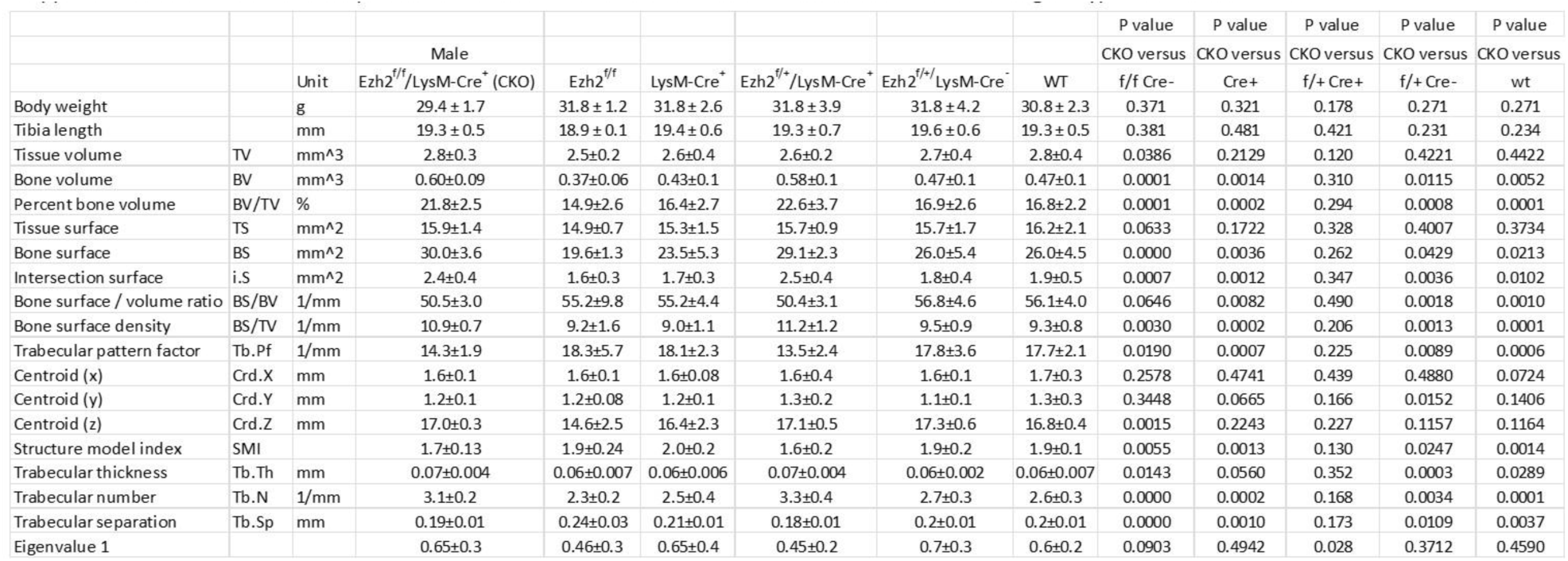
Body weight and tibia length and micro-CT bone parameters of tibia of 6 months old male Ezh2^flox/flox^/LysM-Cre^+^ CKO and other genotypic and wild type control mice. CKO (Ezh2^flox/flox^/LysM-Cre^+^, n=10), Ezh2^f/f^ (Ezh2^flox/flox^, f/f Cre^-^,control, n=8), Cre+ (LysM-Cre^+^ control, n=8), f/+Cre+ (Ezh2^flox/+^/LysM-Cre^+^, n=9), f/+ Cre- (Ezh2^flox/+^/LysM-Cre^-^, n=7) and Wt (wild type control, n=9) male mice. Data are expressed as mean ± SD, p value was analyzed by ttest, CKO versus other genotypic mouse groups.

**Supplemental Table 2.**
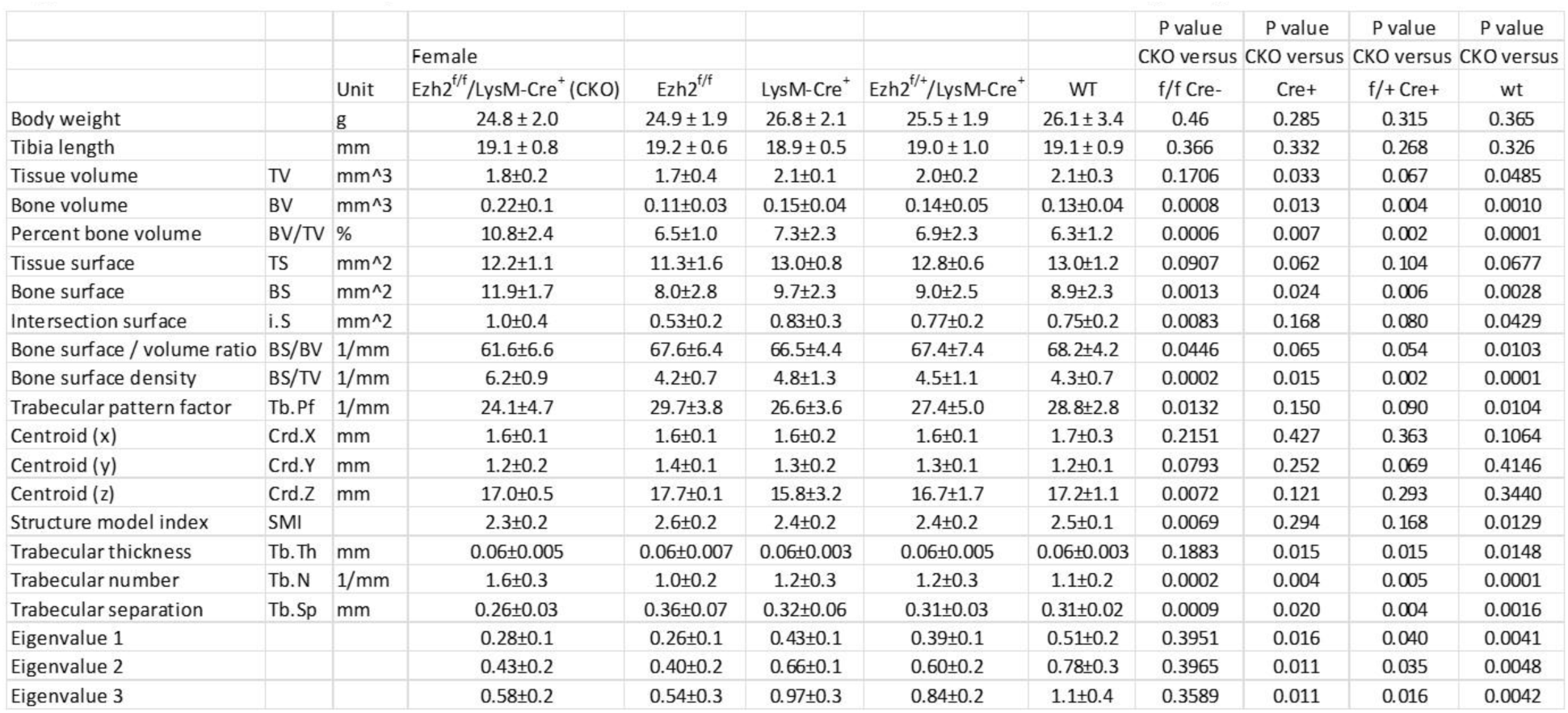
Body weight and tibia length and micro-CT bone parameters of tibia of 6 months old female Ezh2^flox/flox^/LysM-Cre^+^ CKO and other genotypic and wild type control mice. CKO (Ezh2^flox/flox^/LysM-Cre^+^, n=9), Ezh2^f/f^ (Ezh2^flox/flox^, f/f Cre^-^, control, n=8), Cre+ (LysM-Cre^+^ control, n=7), f/+Cre+ (Ezh2^flox/+^/LysM-Cre^+^, n=8), f/+ Cre- (Ezh2^flox/+^/LysM-Cre^-^, n=8) and Wt (wild type control, n=8) male mice. Data are expressed as mean ± SD, p value was analyzed by ttest, CKO versus other genotypic mouse groups.

**Supplemental Table 3.**
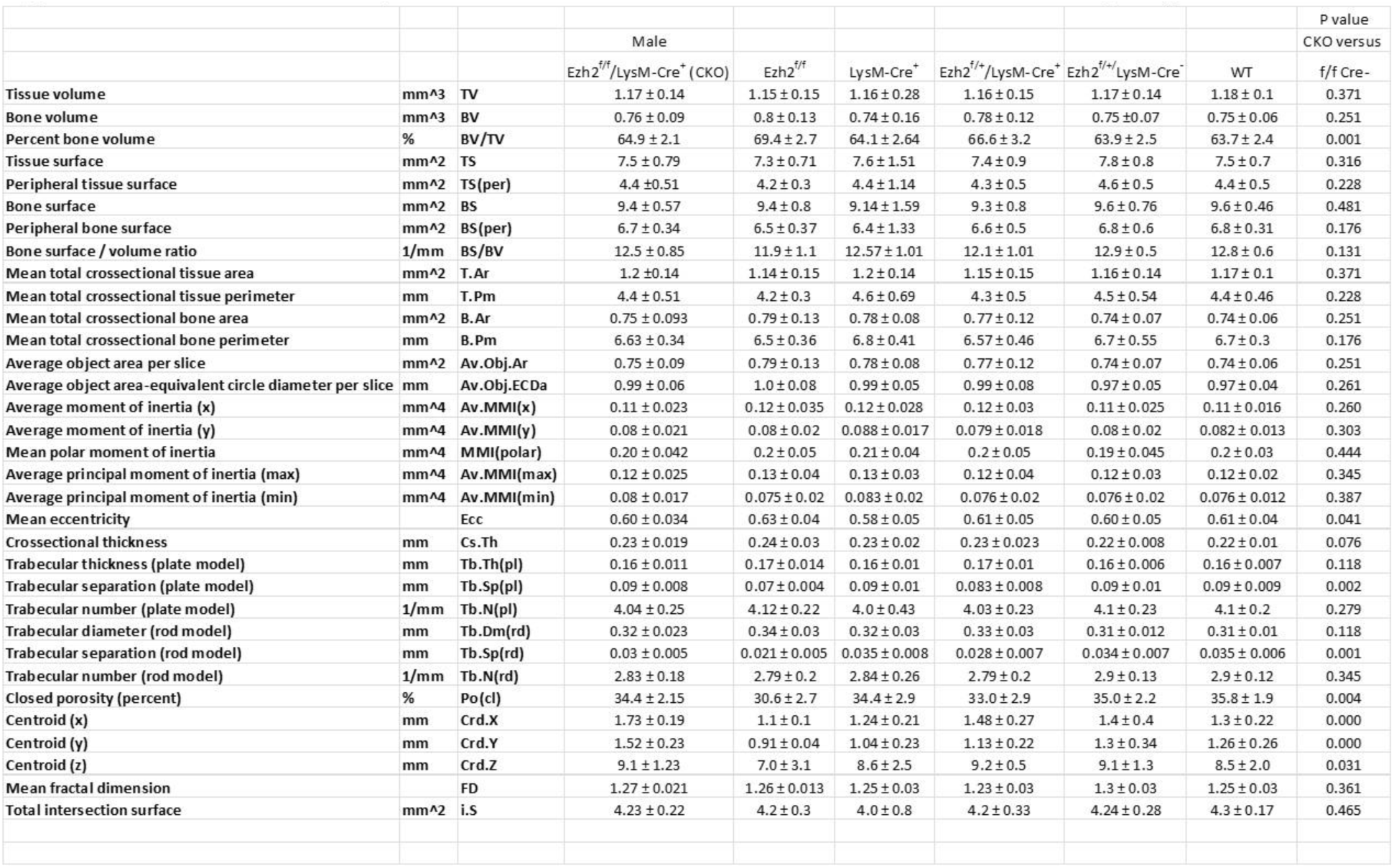
Micro-CT cortical bone parameters of tibia of 6 months old male Ezh2^flox/flox^/LysM-Cre^+^ CKO and other genotypic and wild type control mice. CKO (Ezh2^flox/flox^/LysM-Cre^+^, n=10), Ezh2^f/f^ (Ezh2^flox/flox^, f/f Cre^-^, control, n=8), Cre+ (LysM-Cre^+^ control, n=8), f/+Cre+ (Ezh2^flox/+^/LysM-Cre^+^, n=9), f/+ Cre- (Ezh2^flox/+^/LysM-Cre^-^, n=7) and Wt (wild type control, n=9) male mice. Data are expressed as mean ± SD, p value was analyzed by ttest, CKO versus f/f control.

**Supplemental Table 4.** Micro-CT cortical bone parameters of tibia of 6 months old female Ezh2^flox/flox^/LysM-Cre^+^ CKO and other genotypic and wild type control mice. CKO (Ezh2^flox/flox^/LysM-Cre^+^, n=9), Ezh2^f/f^ (Ezh2^flox/flox^, f/f Cre^-^, control, n=8), Cre+ (LysM-Cre^+^ control, n=7), f/+Cre+ (Ezh2^flox/+^/LysM-Cre^+^, n=8), f/+ Cre- (Ezh2^flox/+^/LysM-Cre^-^, n=8) and Wt (wild type control, n=8) female mice. Data are expressed as mean ± SD, p value was analyzed by ttest, CKO versus other genotypic mouse groups.

|  |  |  | Female |  |  |  |  | P value<br>CKO versus | P value<br>CKO versus | P value<br>CKO versus |
| --- | --- | --- | --- | --- | --- | --- | --- | --- | --- | --- |
|  |  |  | Ezh2<sup>f/f</sup>/LysM-Cre<sup>+</sup> (CKO) | Ezh2<sup>f/f</sup> | LysM-Cre<sup>+</sup> | Ezh2<sup>f/+</sup>/LysM-Cre<sup>+</sup> | WT | f/f Cre<sup>-</sup> | Cre<sup>+</sup> | f/+ Cre<sup>+</sup> |
| Tissue volume | mm<sup>3</sup> | TV | 0.96 ± 0.05 | 0.87 ± 0.05 | 0.95 ± 0.06 | 0.92 ± 0.06 | 0.94 ± 0.07 | 0.001 | 0.385 | 0.054 |
| Bone volume | mm<sup>3</sup> | BV | 0.68 ± 0.05 | 0.64 ± 0.03 | 0.68 ± 0.04 | 0.65 ± 0.02 | 0.66 ± 0.04 | 0.042 | 0.432 | 0.038 |
| Percent bone volume | % | BV/TV | 70.8 ± 4.1 | 73.9 ± 2.2 | 71.1 ± 3.1 | 70.6 ± 2.8 | 70.5 ± 2.7 | 0.055 | 0.444 | 0.438 |
| Tissue surface | mm<sup>2</sup> | TS | 6.9 ± 0.7 | 6.0 ± 0.21 | 6.67 ± 0.76 | 6.4 ± 0.54 | 6.4 ± 0.6 | 0.004 | 0.247 | 0.049 |
| Peripheral tissue surface | mm<sup>2</sup> | TS(per) | 4.25 ± 0.5 | 3.6 ± 0.1 | 4.0 ± 0.54 | 3.9 ± 0.4 | 3.9 ± 0.3 | 0.009 | 0.202 | 0.089 |
| Bone surface | mm<sup>2</sup> | BS | 8.2 ± 0.4 | 7.7 ± 0.3 | 8.2 ± 0.43 | 8.0 ± 0.5 | 8.3 ± 0.7 | 0.013 | 0.377 | 0.181 |
| Peripheral bone surface | mm<sup>2</sup> | BS(per) | 5.77 ± 0.23 | 5.5 ± 0.2 | 5.8 ± 0.2 | 5.7 ± 0.4 | 5.9 ± 0.4 | 0.011 | 0.358 | 0.400 |
| Bone surface / volume ratio | 1/mm | BS/BV | 12.0 ± 0.5 | 12.0 ± 0.45 | 12.1 ± 0.18 | 12.3 ± 0.5 | 12.5 ± 0.6 | 0.462 | 0.264 | 0.102 |
| Mean total crosssectional tissue area | mm<sup>2</sup> | T.Ar | 0.95 ± 0.05 | 0.86 ± 0.05 | 0.95 ± 0.06 | 0.9 ± 0.06 | 0.93 ± 0.07 | 0.001 | 0.385 | 0.054 |
| Mean total crosssectional tissue perimeter | mm | T.Pm | 4.2 ± 0.5 | 3.6 ± 0.1 | 4.0 ± 0.53 | 3.9 ± 0.4 | 3.8 ± 0.3 | 0.009 | 0.202 | 0.089 |
| Mean total crosssectional bone area | mm<sup>2</sup> | B.Ar | 0.67 ± 0.05 | 0.63 ± 0.03 | 0.67 ± 0.04 | 0.64 ± 0.02 | 0.66 ± 0.04 | 0.042 | 0.432 | 0.038 |
| Mean total crosssectional bone perimeter | mm | B.Pm | 5.7 ± 0.23 | 5.4 ± 0.23 | 5.75 ± 0.2 | 5.7 ± 0.4 | 5.8 ± 0.4 | 0.011 | 0.358 | 0.400 |
| Average object area per slice | mm<sup>2</sup> | Av.Obj.Ar | 0.67 ± 0.05 | 0.63 ± 0.03 | 0.67 ± 0.04 | 0.64 ± 0.02 | 0.66 ± 0.04 | 0.042 | 0.432 | 0.038 |
| Average object area-equivalent circle diameter per slice | mm | Av.Obj.ECDa | 0.93 ± 0.03 | 0.9 ± 0.02 | 0.92 ± 0.03 | 0.9 ± 0.02 | 0.91 ± 0.03 | 0.041 | 0.437 | 0.038 |
| Average moment of inertia (x) | mm<sup>4</sup> | Av.MMI(x) | 0.078 ± 0.01 | 0.064 ± 0.01 | 0.08 ± 0.01 | 0.07 ± 0.007 | 0.077 ± 0.01 | 0.014 | 0.398 | 0.029 |
| Average moment of inertia (y) | mm<sup>4</sup> | Av.MMI(y) | 0.053 ± 0.007 | 0.05 ± 0.006 | 0.053 ± 0.005 | 0.05 ± 0.006 | 0.055 ± 0.008 | 0.171 | 0.492 | 0.309 |
| Mean polar moment of inertia | mm<sup>4</sup> | MMI(polar) | 0.13 ± 0.02 | 0.11 ± 0.01 | 0.13 ± 0.01 | 0.12 ± 0.01 | 0.13 ± 0.02 | 0.021 | 0.430 | 0.081 |
| Average principal moment of inertia (max) | mm<sup>4</sup> | Av.MMI(max) | 0.08 ± 0.01 | 0.068 ± 0.007 | 0.08 ± 0.01 | 0.07 ± 0.008 | 0.079 ± 0.015 | 0.018 | 0.444 | 0.031 |
| Average principal moment of inertia (min) | mm<sup>4</sup> | Av.MMI(min) | 0.052 ± 0.007 | 0.046 ± 0.006 | 0.05 ± 0.005 | 0.05 ± 0.006 | 0.053 ± 0.006 | 0.037 | 0.414 | 0.322 |
| Mean eccentricity |  | Ecc | 0.59 ± 0.05 | 0.58 ± 0.02 | 0.58 ± 0.04 | 0.53 ± 0.04 | 0.57 ± 0.05 | 0.310 | 0.457 | 0.006 |
| Crosssectional thickness | mm | Cs.Th | 0.24 ± 0.01 | 0.23 ± 0.009 | 0.23 ± 0.008 | 0.23 ± 0.01 | 0.23 ± 0.01 | 0.400 | 0.302 | 0.043 |
| Trabecular thickness (plate model) | mm | Tb.Th(pl) | 0.17 ± 0.007 | 0.17 ± 0.006 | 0.16 ± 0.002 | 0.16 ± 0.006 | 0.16 ± 0.008 | 0.450 | 0.238 | 0.097 |
| Trabecular separation (plate model) | mm | Tb.Sp(pl) | 0.069 ± 0.01 | 0.059 ± 0.006 | 0.067 ± 0.01 | 0.068 ± 0.007 | 0.067 ± 0.006 | 0.034 | 0.374 | 0.405 |
| Trabecular number (plate model) | 1/mm | Tb.N(pl) | 4.25 ± 0.18 | 4.44 ± 0.13 | 4.3 ± 0.16 | 4.3 ± 0.1 | 4.4 ± 0.13 | 0.017 | 0.210 | 0.098 |
| Trabecular diameter (rod model) | mm | Tb.Dm(rd) | 0.33 ± 0.02 | 0.33 ± 0.01 | 0.33 ± 0.005 | 0.33 ± 0.01 | 0.32 ± 0.016 | 0.450 | 0.238 | 0.097 |
| Trabecular separation (rod model) | mm | Tb.Sp(rd) | 0.018 ± 0.01 | 0.010 ± 0.005 | 0.017 ± 0.008 | 0.018 ± 0.006 | 0.018 ± 0.006 | 0.049 | 0.425 | 0.493 |
| Trabecular number (rod model) | 1/mm | Tb.N(rd) | 2.85 ± 0.1 | 2.9 ± 0.09 | 2.9 ± 0.05 | 2.9 ± 0.07 | 2.95 ± 0.1 | 0.082 | 0.163 | 0.048 |
| Closed porosity (percent) | % | Po(cl) | 26.8 ± 2.7 | 26.0 ± 2.2 | 27.9 ± 1.0 | 28.5 ± 2.4 | 29.3 ± 2.7 | 0.284 | 0.159 | 0.087 |
| Centroid (x) | mm | Crđ.X | 1.68 ± 0.25 | 1.2 ± 0.1 | 1.5 ± 0.08 | 1.5 ± 0.3 | 1.15 ± 0.15 | 0.000 | 0.062 | 0.049 |
| Centroid (y) | mm | Crđ.Y | 1.35 ± 0.16 | 1.3 ± 0.29 | 1.1 ± 0.14 | 1.24 ± 0.25 | 0.97 ± 0.17 | 0.495 | 0.005 | 0.130 |
| Centroid (z) | mm | Crđ.Z | 8.9 ± 0.5 | 9.4 ± 0.31 | 7.45 ± 3.1 | 8.9 ± 1.3 | 9.0 ± 0.9 | 0.019 | 0.074 | 0.472 |
| Mean fractal dimension |  | FD | 1.2 ± 0.03 | 1.24 ± 0.03 | 1.23 ± 0.02 | 1.24 ± 0.03 | 1.24 ± 0.02 | 0.387 | 0.336 | 0.429 |
| Total intersection surface | mm<sup>2</sup> | I.S | 3.8 ± 0.12 | 3.62 ± 0.1 | 3.78 ± 0.12 | 3.69 ± 0.1 | 3.8 ± 0.27 | 0.011 | 0.432 | 0.063 |

**Supplemental Table 5.**
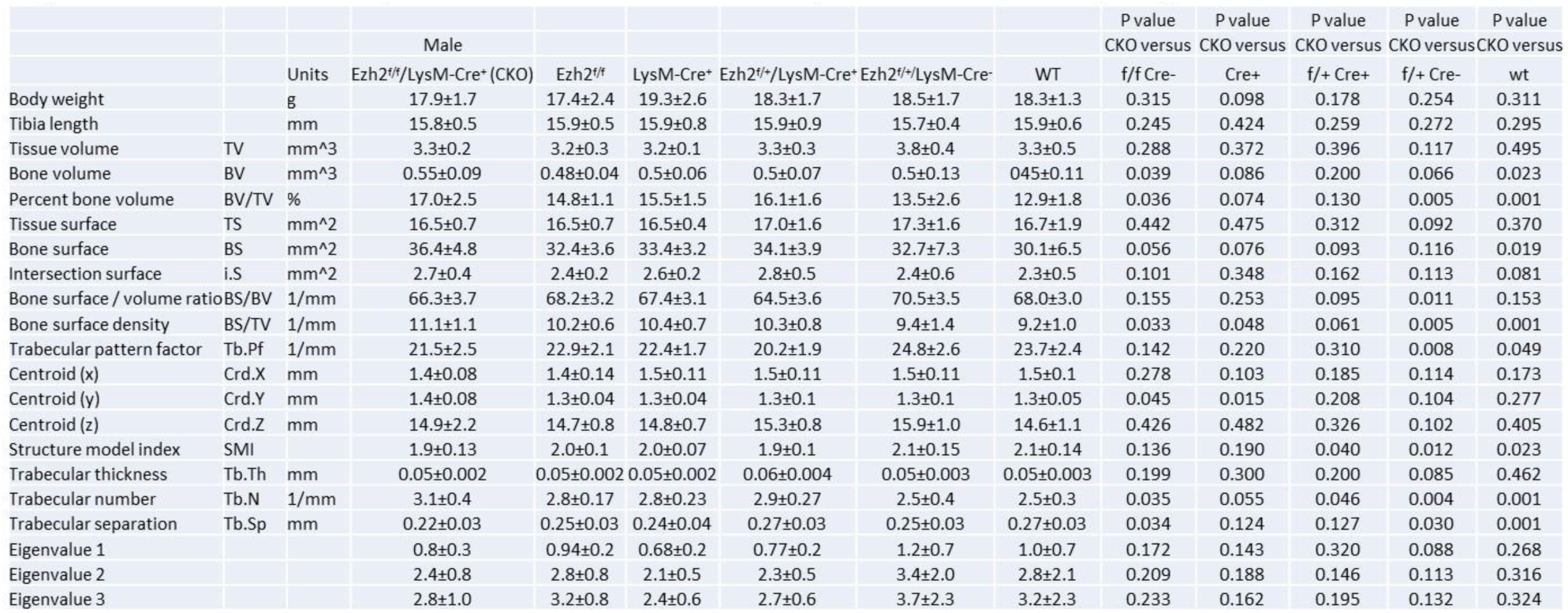
Body weight and tibia length and micro-CT trabecular bone parameters of tibia of 30 days old male Ezh2^flox/flox^/LysM-Cre^+^ CKO and other genotypic and wild type control mice. CKO (Ezh2^flox/flox^/LysM-Cre^+^, n=10), Ezh2^f/f^ (Ezh2^flox/flox^, f/f Cre^-^, control, n=8), Cre+ (LysM-Cre^+^ control, n=8), f/+Cre+ (Ezh2^flox/+^/LysM-Cre^+^, n=9), f/+ Cre- (Ezh2^flox/+^/LysM-Cre^-^, n=7) and Wt (wild type control, n=9) male mice. Data are expressed as mean ± SD, p value was analyzed by ttest, CKO versus other genotypic mouse groups.

**Supplemental Table 6.**
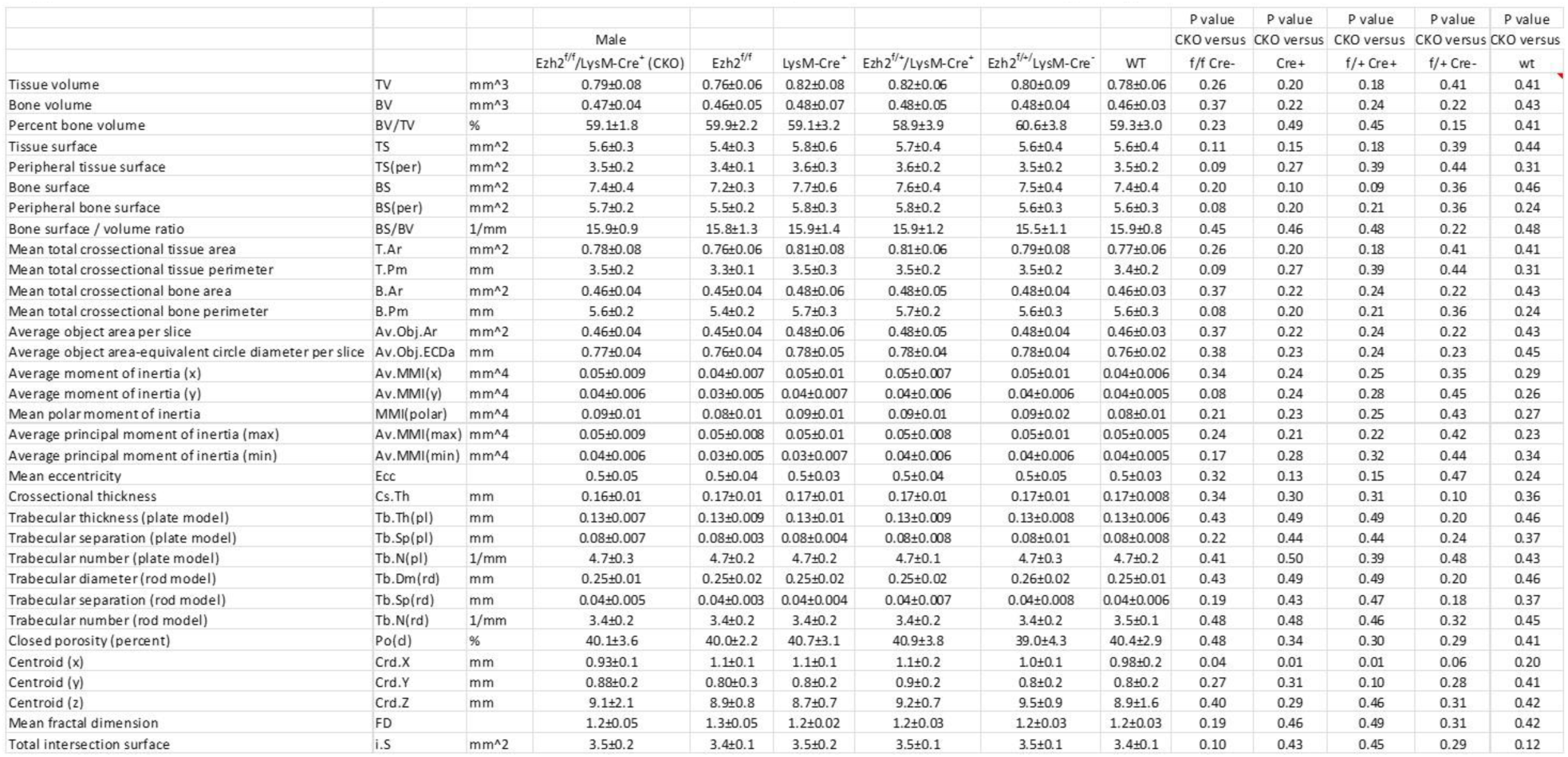
Micro-CT cortical bone parameters of tibia of 30 days old male Ezh2^flox/flox^/LysM-Cre^+^ CKO and other genotypic and wild type control mice. CKO (Ezh2^flox/flox^/LysM-Cre^+^, n=10), Ezh2^f/f^ (Ezh2^flox/flox^, f/f Cre^-^, control, n=8), Cre+ (LysM-Cre^+^ control, n=8), f/+Cre+ (Ezh2^flox/+^/LysM-Cre^+^, n=9), f/+ Cre- (Ezh2^flox/+^/LysM-Cre^-^, n=7) and Wt (wild type control, n=9) male mice. Data are expressed as mean ± SD, p value was analyzed by ttest, CKO versus other genotypic and control mouse groups.

**Supplemental Table 7.**
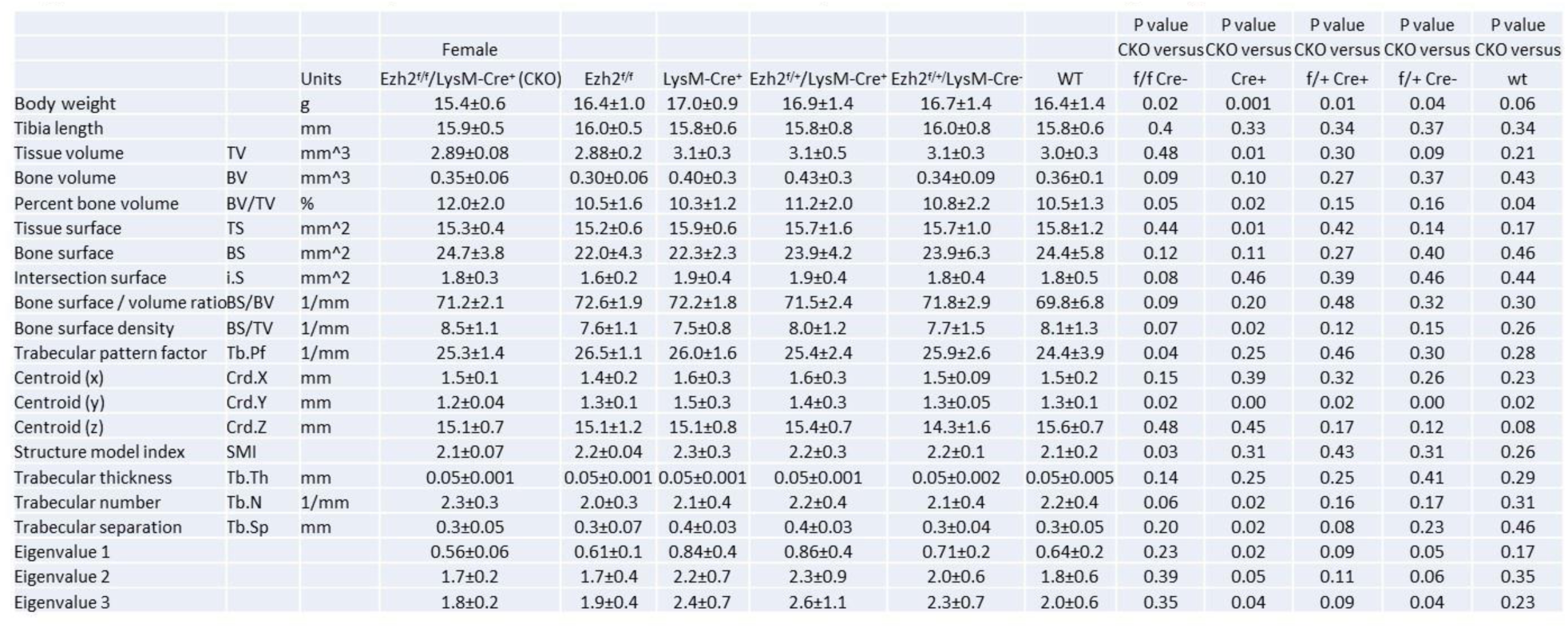
Body weight and tibia length and micro-CT trabecular bone parameters of tibia of 30 days old female Ezh2^flox/flox^/LysM-Cre^+^ CKO and other genotypic and wild type control mice. CKO (Ezh2^flox/flox^/LysM-Cre^+^, n=9), Ezh2^f/f^ (Ezh2^flox/flox^, f/f Cre^-^, control, n=8), Cre+ (LysM-Cre^+^ control, n=7), f/+Cre+ (Ezh2^flox/+^/LysM-Cre^+^, n=8), f/+ Cre- (Ezh2^flox/+^/LysM-Cre^-^, n=8) and Wt (wild type control, n=8) female mice. Data are expressed as mean ± SD, p value was analyzed by ttest, CKO versus other genotypic mouse groups.

**Supplemental Table 8.**
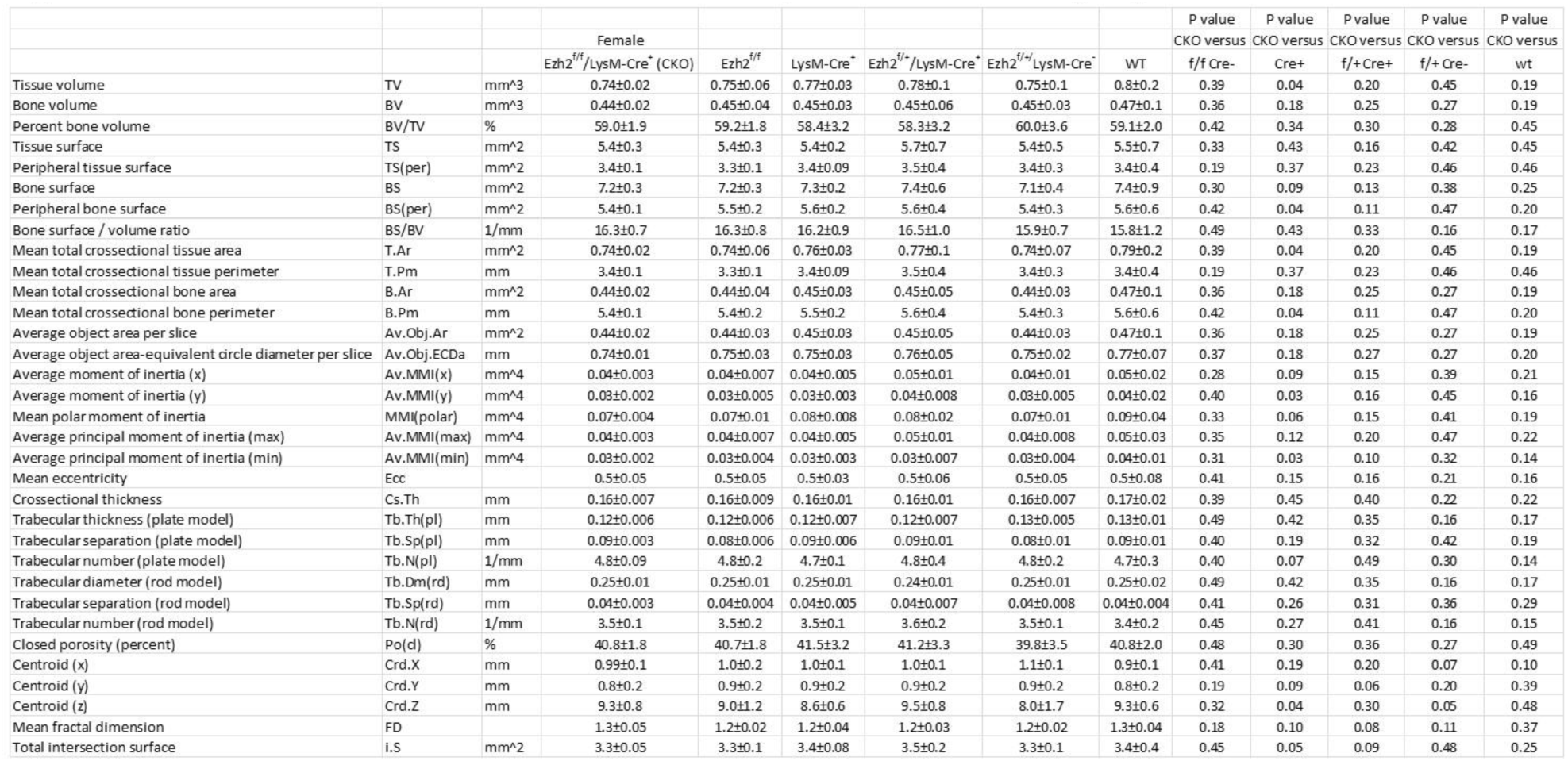
Micro-CT cortical bone parameters of tibia of 30 days old female Ezh2^flox/flox^/LysM-Cre^+^ CKO and other genotypic and wild type control mice. CKO (Ezh2^flox/flox^/LysM-Cre^+^, n=10), Ezh2^f/f^ (Ezh2^flox/flox^, f/f Cre^-^, control, n=8), Cre+ (LysM-Cre^+^ control, n=8), f/+Cre+ (Ezh2^flox/+^/LysM-Cre^+^, n=9), f/+ Cre- (Ezh2^flox/+^/LysM-Cre^-^, n=7) and Wt (wild type control, n=9) female mice. Data are expressed as mean ± SD, p value was analyzed by ttest, CKO versus other genotypic and control mouse groups.

**Supplemental Table 9.**
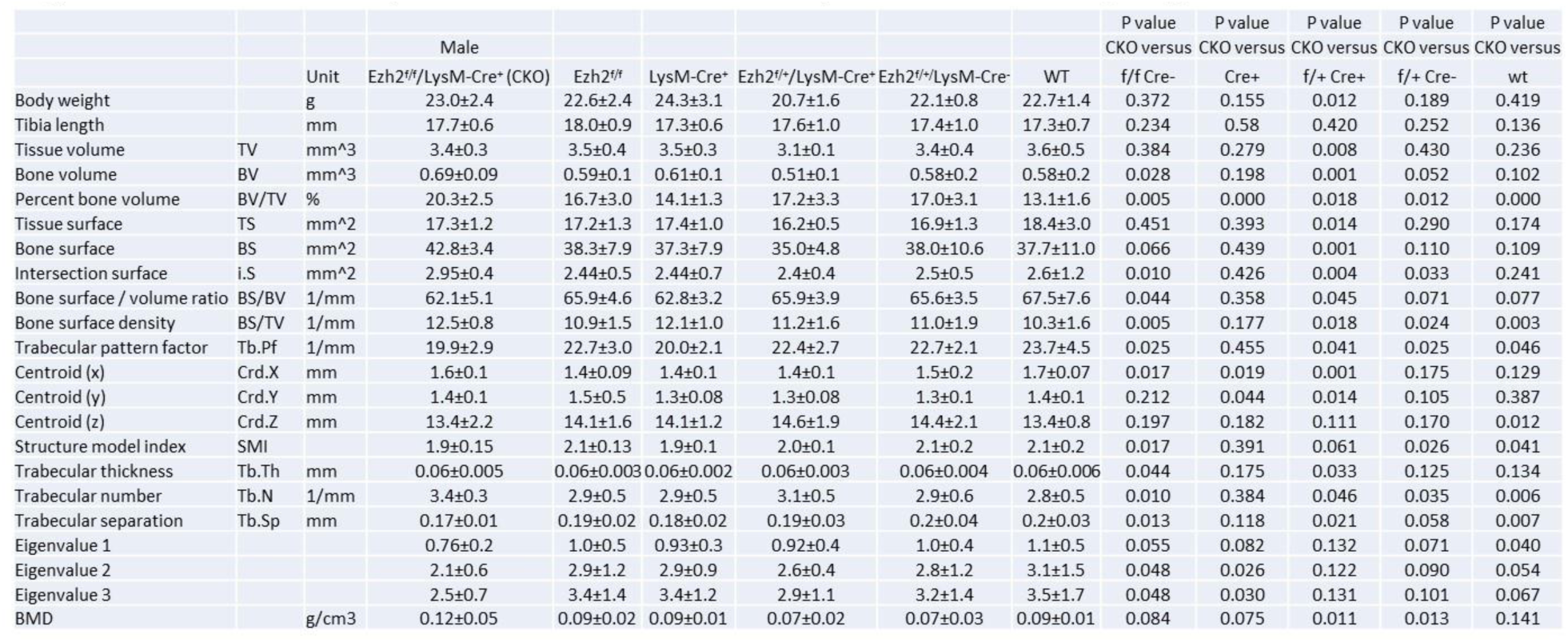
Body weight and tibia length and micro-CT trabecular bone parameters of tibia of 56 days old male Ezh2^flox/flox^/LysM-Cre^+^ CKO and other genotypic and wild type control mice. CKO (Ezh2^flox/flox^/LysM-Cre^+^, n=9), Ezh2^f/f^ (Ezh2^flox/flox^, f/f Cre^-^, control, n=8), Cre+ (LysM-Cre^+^ control, n=7), f/+Cre+ (Ezh2^flox/+^/LysM-Cre^+^, n=8), f/+ Cre- (Ezh2^flox/+^/LysM-Cre^-^, n=8) and Wt (wild type control, n=8) male mice. Data are expressed as mean ± SD, p value was analyzed by ttest, CKO versus other genotypic mouse groups.

**Supplemental Table 10.**
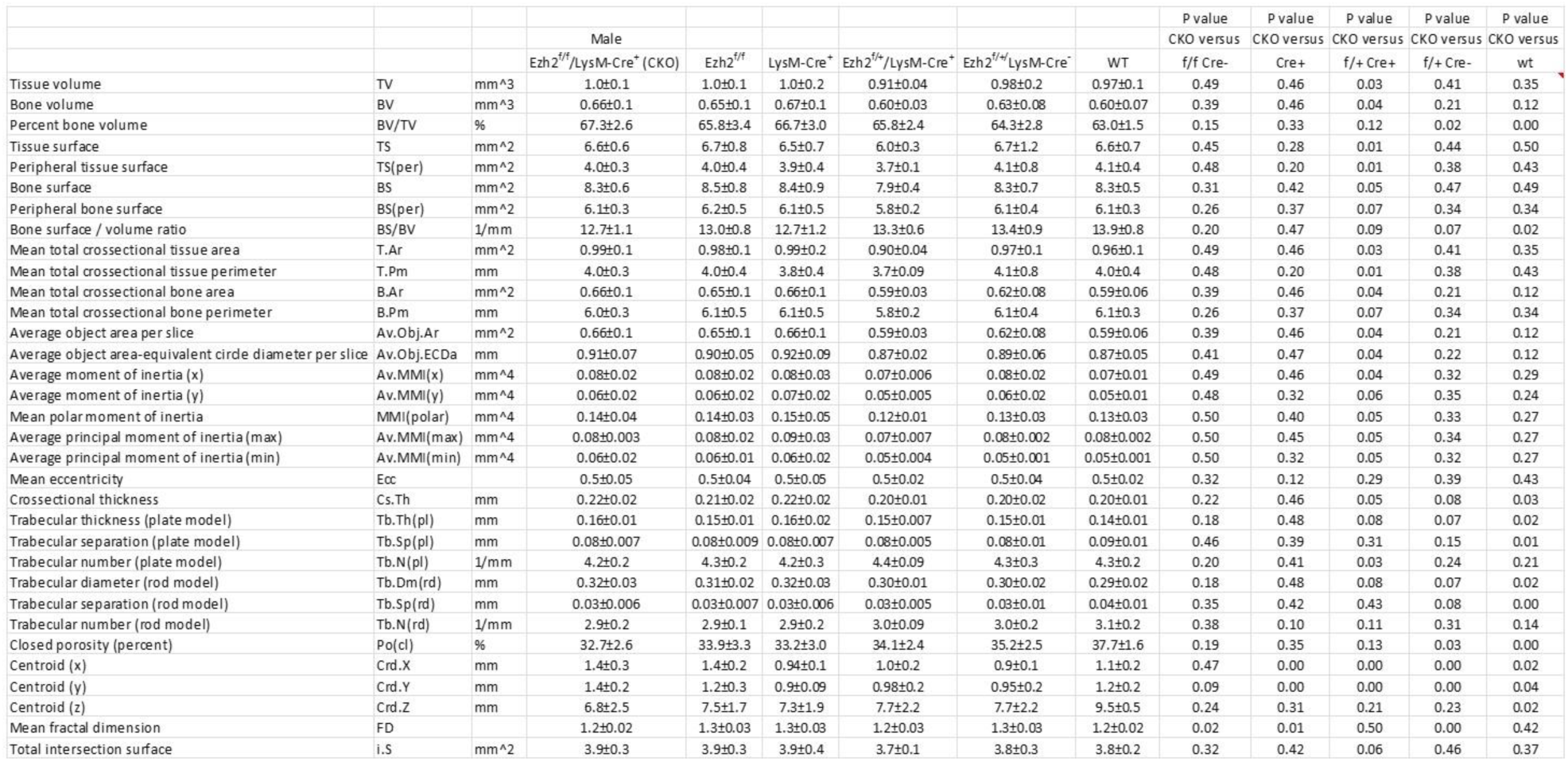
Micro-CT cortical bone parameters of tibia of 56 days old male Ezh2^flox/flox^/LysM-Cre^+^ CKO and other genotypic and wild type control mice. CKO (Ezh2^flox/flox^/LysM-Cre^+^, n=10), Ezh2^f/f^ (Ezh2^flox/flox^, f/f Cre^-^, control, n=8), Cre+ (LysM-Cre^+^ control, n=8), f/+Cre+ (Ezh2^flox/+^/LysM-Cre^+^, n=9), f/+ Cre- (Ezh2^flox/+^/LysM-Cre^-^, n=7) and Wt (wild type control, n=9) male mice. Data are expressed as mean ± SD, p value was analyzed by ttest, CKO versus other genotypic and control mouse groups.

**Supplemental Table 11.**
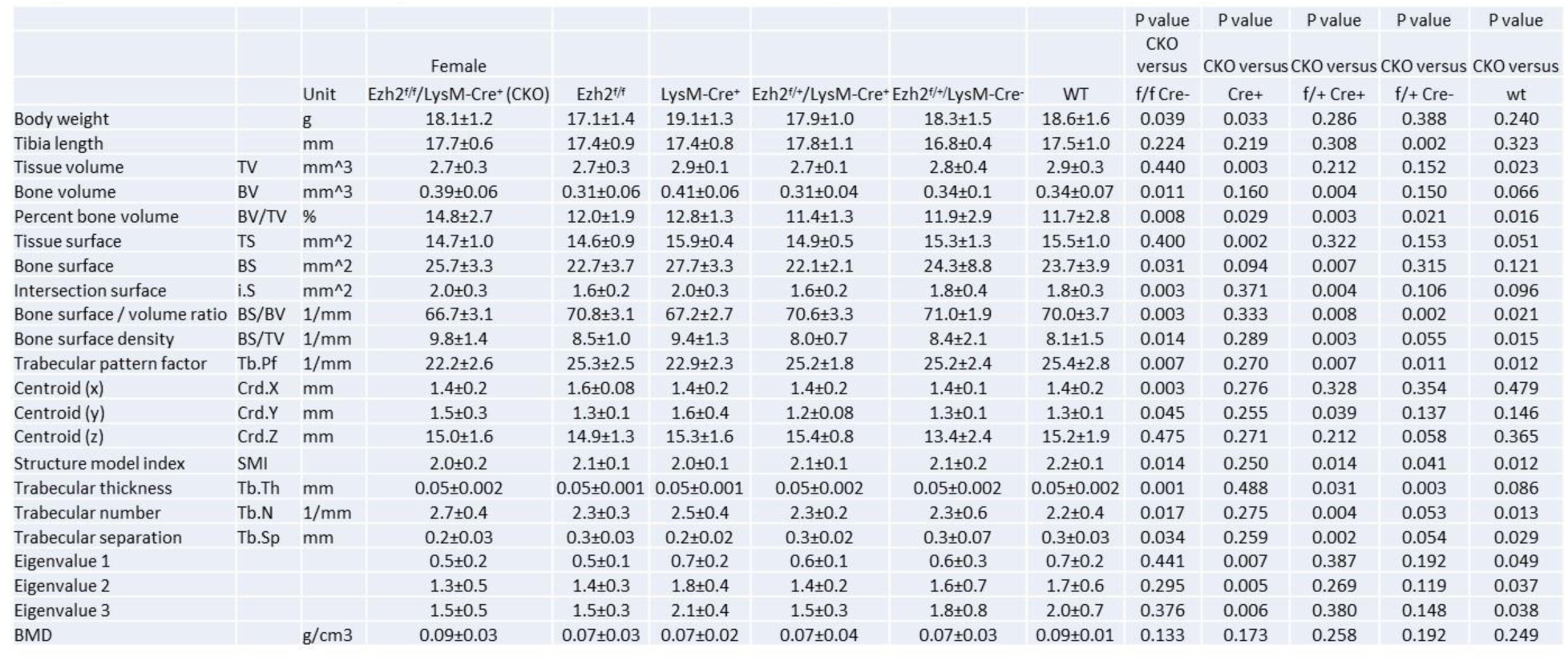
Body weight and tibia length and micro-CT trabecular bone parameters of tibia of 56 days old female Ezh2^flox/flox^/LysM-Cre^+^ CKO and other genotypic and wild type control mice. CKO (Ezh2^flox/flox^/LysM-Cre^+^, n=9), Ezh2^f/f^ (Ezh2^flox/flox^, f/f Cre^-^, control, n=8), Cre+ (LysM-Cre^+^ control, n=7), f/+Cre+ (Ezh2^flox/+^/LysM-Cre^+^, n=8), f/+ Cre- (Ezh2^flox/+^/LysM-Cre^-^, n=8) and Wt (wild type control, n=8) female mice. Data are expressed as mean ± SD, p value was analyzed by ttest, CKO versus other genotypic mouse groups.

**Supplemental Table 12.**
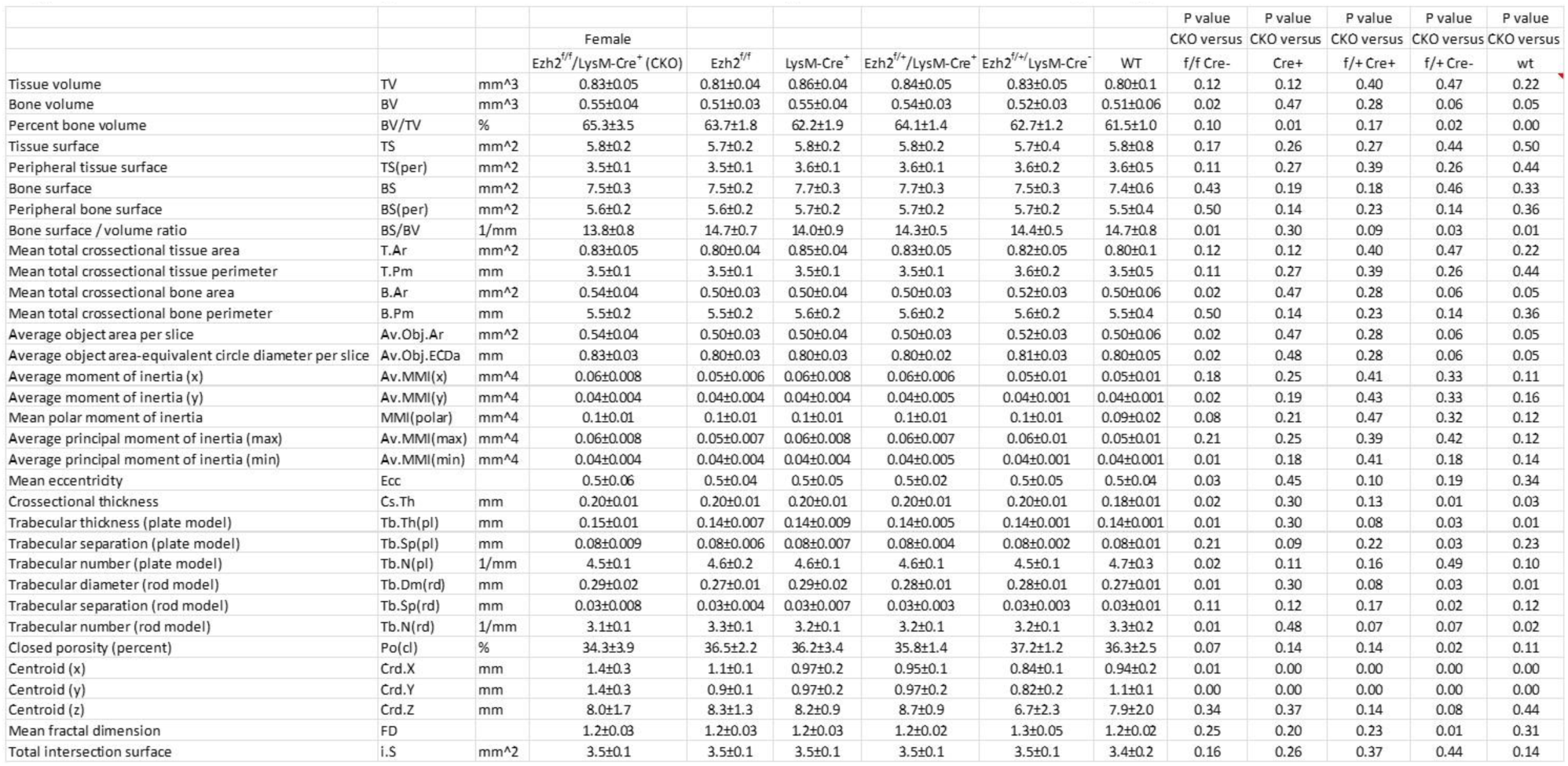
Micro-CT cortical bone parameters of tibia of 56 days old female Ezh2^flox/flox^/LysM-Cre^+^ CKO and other genotypic and wild type control mice. CKO (Ezh2^flox/flox^/LysM-Cre^+^, n=10), Ezh2^f/f^ (Ezh2^flox/flox^, f/f Cre^-^, control, n=8), Cre+ (LysM-Cre^+^ control, n=8), f/+Cre+ (Ezh2^flox/+^/LysM-Cre^+^, n=9), f/+ Cre- (Ezh2^flox/+^/LysM-Cre^-^, n=7) and Wt (wild type control, n=9) female mice. Data are expressed as mean ± SD, p value was analyzed by ttest, CKO versus other genotypic and control mouse groups.

